# Biodiversity turnover in coastal marine eukaryotic plankton revealed by a 20-year metabarcoding time series

**DOI:** 10.64898/2026.11.25.768829

**Authors:** Joanna Warwick-Dugdale, Clare Ostle, Kimberley Bird, Yousef Touhami, Rohan Allen, Cordelia Roberts, Laura Branscombe, Katherine Helliwell, Nicolas Henry, Nicolas Djeghri, Tim Smyth, Michael Cunliffe

**Affiliations:** Marine Biological Association, The Laboratory, Citadel Hill, Plymouth, UK; School of Biological and Marine Sciences, University of Plymouth, Plymouth, UK; School of Biosciences, University of Exeter, Exeter, UK; CNRS, Sorbonne Université, FR2424, ABiMS, Station Biologique de Roscoff, Roscoff, France; Research Federation for the Study of Global Ocean Systems Ecology and Evolution, FR2022/GOSEE, Paris 75016, France; Laboratory of Environmental Marine Sciences, University of Western Brittany, 29200 Brest, France; Plymouth Marine Laboratory, Prospect Place, Plymouth, UK

**Keywords:** marine plankton, metabarcoding, time series, regime shift, biodiversity turnover, North Atlantic Oscillation, coastal ecosystem, eukaryotic microbes

## Abstract

Long-term observations are essential to resolve how ecosystems respond to environmental variability, yet decadal-scale molecular time series of marine plankton communities remain rare. Here, we analysed a >20-year (2001–2023) high-frequency 18S rRNA gene (V9) metabarcoding time series of eukaryotic plankton from a coastal observatory in the Western English Channel. Community composition exhibited recurrent seasonal cycles driven primarily by seawater temperature and water column stratification. Superimposed on this seasonal baseline, we identified a pronounced but transient community shift between 2008 and 2011, characterised by reduced diversity, and distinct community structuring, indicating plankton ecosystem-wide biodiversity turnover. This period reflected marked community reorganisation, with potential declines in larger plankton taxa (including copepods and diatoms) and concurrent increases in picoeukaryotes and heterotrophic stramenopiles, consistent with a shift toward smaller-sized, microbial communities. While temperature explained much of the seasonal-scale variation, it did not account for the proposed biodiversity turnover. Instead, the shift was associated with negative phases of the North Atlantic Oscillation, implicating basin-scale climate forcing. Following 2011, communities returned to a state resembling earlier years, indicating a reversible, rather than persistent, change. These findings demonstrate that coastal plankton communities can undergo rapid, climate-driven but reversible biodiversity turnover that is not captured by seasonal dynamics alone and highlight the power of long-term metabarcoding time series to detect transient and recoverable ecologically significant community reorganisation.

## Introduction

Marine plankton communities respond rapidly to environmental variability, making them sensitive indicators of ecosystem responses to change. Temperature, nutrient availability, water-column structure and large-scale atmospheric processes all can regulate plankton phenology, diversity and community composition, with consequences for food webs and biogeochemical cycling (Richardson & Schoeman, 2004; Doney *et al*., 2012). Long-term observations have revealed substantial reorganisations of plankton communities, including shifts in North Atlantic copepod biogeography and changes propagating across trophic levels (Beaugrand *et al*., 2002; Richardson & Schoeman, 2004). Historical reconstructions further indicate that contemporary plankton assemblages exhibit progressive divergence from their pre-industrial states (Jonkers *et al*., 2019).

A major challenge is to distinguish recurrent ecological variability from more consequential biodiversity change. Temperate plankton communities undergo pronounced seasonal succession and can return to broadly similar community configurations each year despite substantial environmental variability (Gilbert *et al*., 2009, 2012; Giovannoni & Vergin, 2012; Lambert *et al*., 2019). Superimposed upon this seasonal succession, environmental change can drive substantial reorganisation of community composition. Recently, temporal turnover (i.e. the replacement or redistribution of taxa through time) has been recognised as a major dimension of biodiversity change, with accelerated compositional turnover linked to temperature variation (Pinsky *et al*., 2025). At its most pronounced, community reorganisation may resemble an ecological regime shift. Yet such changes need not be permanent, instead strong environmental forcing may produce transient biodiversity turnover followed by recovery towards an earlier community state (Pinsky *et al*., 2025). Distinguishing persistent from reversible change is therefore central to understanding ecological resilience but requires sufficiently long observations to capture both community reorganisation and potential recovery.

Long-term plankton observations have been fundamental in detecting climate-associated ecological change (Beaugrand *et al*., 2002; Richardson & Schoeman, 2004). However, many historical records rely principally on microscopy, abundance or biomass measurements and consequently provide limited resolution of small, cryptic, and morphologically indistinct eukaryotes. Environmental DNA (eDNA) metabarcoding provides a complementary perspective, simultaneously characterising organisms across broad phylogenetic and size ranges. The power of this approach has been demonstrated where metabarcoding has revealed extensive and previously unresolved eukaryotic diversity across the global ocean (de Vargas *et al*., 2015; Sunagawa *et al*., 2020). In contrast to this increasingly detailed understanding of spatial molecular diversity, comparable temporal resolution remains limited. Molecular time series have demonstrated strong seasonal recurrence in marine microbial communities (Gilbert *et al*., 2009, 2012; Ward *et al*., 2017; Lambert *et al*., 2019), but few span the decades required to distinguish seasonal recurrence from long-term change, episodic turnover and subsequent recovery.

Large-scale climate variability provides an important context for examining plankton dynamics in the North Atlantic. The North Atlantic Oscillation (NAO), a dominant mode of atmospheric variability in the region, affects marine ecosystems through changes in winds, temperature, circulation, vertical mixing, and nutrient supply. NAO variability has been associated with changes in copepod abundance and distribution (Fromentin & Planque, 1996; Beaugrand *et al*., 2002), English Channel spring phytoplankton composition (Irigoien *et al*., 2000), and North Atlantic phytoplankton community structure and carbon flux (Henson *et al*., 2012). Whether such climatic variability is associated with coordinated turnover across the broader eukaryotic plankton community, including microbial groups poorly represented by conventional monitoring, remains unclear.

The Western Channel Observatory (WCO) Station L4 in the Western English Channel provides an exceptional system in which to address this knowledge gap. Station L4 in the WCO is a seasonally stratified coastal site supported by long-term physical, chemical and biological observations (McEvoy *et al*., 2023). Previous eDNA metabarcoding studies have demonstrated recurrent seasonal structuring of bacterioplankton and fungal communities at Station L4 (Gilbert *et al*., 2009, 2012; Chrismas *et al*, 2023), while sustained sampling has generated an extensive eDNA archive demonstrating its potential for reconstructing ecological dynamics over timescales rarely achieved using molecular approaches.

Here, we use a >20-year (2001–2023), high-frequency 18S rRNA V9 metabarcoding time series of the eukaryotic plankton community at Station L4. By combining approximately weekly molecular observations with long-term environmental data and a range of complementary analytical approaches we ask: (i) how consistently does plankton biodiversity recur seasonally across two decades; (ii) is this seasonal template interrupted by periods of pronounced biodiversity turnover; (iii) if turnover occurs, does it represent persistent change or transient reorganisation followed by recovery; and (iv) how are these dynamics associated with local hydrographic conditions and basin-scale climate variability, particularly the NAO? This long-term molecular perspective enables recurrent seasonal dynamics to be separated from episodic community reorganisation and provides a framework for assessing the resilience of coastal plankton biodiversity to climate variability.

## Methods

### Collection of samples

This study combines time series eDNA samples based on three campaigns, collected between February 2001 and February 2023. Samples collected between February 2002 and December 2018 were previously surveyed for fungal diversity using the ‘Fungi-specific’ internal transcribed spacer (ITS) region as reported in Chrismas *et al* (2023). Here, the entire eDNA library time series (2001-2023) were sequenced and processed in a continuous campaign to characterise the eukaryotic plankton community as detailed below.

Samples were collected approximately once per week from the surface (≤5 m depth) of the Western English Channel (WEC) coastal Western Channel Observatory (WCO) site ‘L4’ (50∘15.00′ N, 4∘13.02′ W; site depth ∼55 m). From the years 2001-2019, seawater was collected via Niskin-rosette from RV Plymouth Quest. 1 L was filtered onto 0.45 µm cellulose-nitrate (516 samples) or polycarbonate filters (46 samples distributed across the following years: 2010 = 1; 2012 = 5; 2013 = 15; 2015 = 7; 2016 = 1; 2018 = 11; 2019 = 6) and stored at −80°C (Chrismas *et al,* 2023). Samples obtained in the years 2022-2023 were collected from the RV MBA Sepia, either via Niskin bottle (2 L volume filtered per sample) or by the under-way sampling system (1 L volume filtered per sample). These samples were collected onto 0.22 µm polyethersulfone filters (13 samples) or cellulose-nitrate filters (25 samples) and stored with nucleic acid preservative (DNA/RNA Sheild, Zymo Research) at −80°C and -20°C (respectively).

### Environmental data

Corresponding environmental parameters were collected by WCO operations (https://www.westernchannelobservatory.org.uk), including temperature, salinity and nutrients (ammonia, nitrate, nitrite-nitrate, phosphate; silicate). Riverine input at time of sampling was calculated as the Total Gauged Daily Flow (GDF; m^3^/s), measured from six sites where water courses run into the Plymouth Sound (National River Flow Archive) (locations illustrated in Chrismas *et al,* 2023; Figure1.a). Monthly mean North Atlantic Oscillation (NAO) index data was downloaded from the National Oceanic and Atmospheric Administration (NOAA: https://www.cpc.ncep.noaa.gov/products/precip/CWlink/pna/norm.nao.monthly.b5001.current.ascii.table; accessed: 14/03/2025).

### DNA extraction, metabarcode amplification and sequencing

The majority of samples were extracted with the ZymoBIOMICS DNA Miniprep Kit (Zymo Research) with 12 samples extracted with the ZymoBIOMICS DNA/RNA Miniprep Kit (Zymo Research). Filters were dissected using a sterile scalpel. Filter sections were added to 700 µl of lysis solution in lysis tubes (ZymoBIOMICS kit components), and were bead beaten three times for one minute duration at 10 m·s^-1^, with five-minute intervals on ice. From here, DNA extraction followed manufacturer’s protocols with 100 µl final elution volume in DNase/RNase Free Water. Possible kit/extraction associated contaminants were determined via the inclusion of one blank (i.e. no sample) sample per extraction batch of ∼20 samples. The ‘Total-eukaryote’ 18S-V9 region rRNA gene was amplified using PCR by NU-OMICS at Northumbria University, with the forward primer 1391f (5’-GTACACACCGCCCGTC -3’) and the reverse primer EukBr (5’-TGATCCTTCTGCAGGTTCACCTAC -3’) (based on Amaral-Zettler *et al*, 2009, and Stoeck *et al*, 2010), according to the Earth Microbiome Project (EMP) protocol (including reference to Caporaso *et al.,* 2012), excepting the use here of KAPA2G Robust Master Mix (Roche Diagnostics). Briefly, the PCR reaction mixture (total volume 25.0 µl) consisted of the following: 13.0 µl of PCR-grade water; 10.0 µl PCR master mix; 0.5 µl of forward primer (10 µM); 0.5 µl of reverse primer (10 µM); 1 µl of template DNA. Samples were amplified in triplicate using the following conditions: Initial denaturing at 94°C for 180 seconds, followed by 35 cycles of: 94°C for 45 seconds, 57°C for 60 seconds, and 72°C for 90 seconds, and then a final elongation step at 72°C for 600 seconds. After assaying size and quantity (using Agilent BioAnalyzer 2100 with High Sensitivity DNA kit), triplicate PCR reactions were combined, cleaned (with 1x Ampure XP beads as per manufacturer’s protocol) and purity was ascertained (i.e. A260/A280 ratio, via nanodrop analysis). Amplicons were sequenced on the Illumina MiSeq platform by NU-OMICS at Northumbria University as 250 bp paired-end reads.

### Sequence processing

The bioinformatic pipeline is summarised in Supporting Figure 1. In total, 20,941,365 18S-V9 rRNA reads were generated from 575 samples (plus 32 extraction kit blanks and 20 sequencing negative controls). Reads were demultiplexed and sequencing adapters removed on the Illumina instrument using GenerateFASTQWorker v2.5.56.9, prior to trimming of primers with CutAdapt v1.5 (Martin, 2011). Amplicon sequences were further processed using nf-core/ampliseq v2.8.0 (doi: 10.5281/zenodo.1493841) (Straub *et al*., 2020) of the nf-core collection of workflows (Ewels *et al*., 2020), utilising reproducible software environments from the Bioconda (Grüning *et al*., 2018) and Biocontainers (da Veiga Leprevost *et al*., 2017) projects. Data quality was evaluated with FastQC v0.12.1 (Andrews, 2010, https://www.bioinformatics.babraham.ac.uk/projects/fastqc/) and summarised with MultiQC (Ewels *et al*., 2016). Sequences were processed independently, then re-evaluated as one pool (pseudo-pooled) with DADA2 (Callahan *et al*., 2016) for additional trimming (including elimination of PhiX contamination, trimming of forward reads at 80 bp and reverse reads at 80 bp (reads shorter than 80 bp were discarded)), removal of low-quality reads (discarding reads with > 3 expected errors), error correction, merging of read pairs, removal of PCR chimeras, and determination and dereplication of individual biological sequences (i.e. Amplicon Sequence Variants: ASVs; total 8860). An average of 19,206 reads were retained per sample.

**Figure 1.**
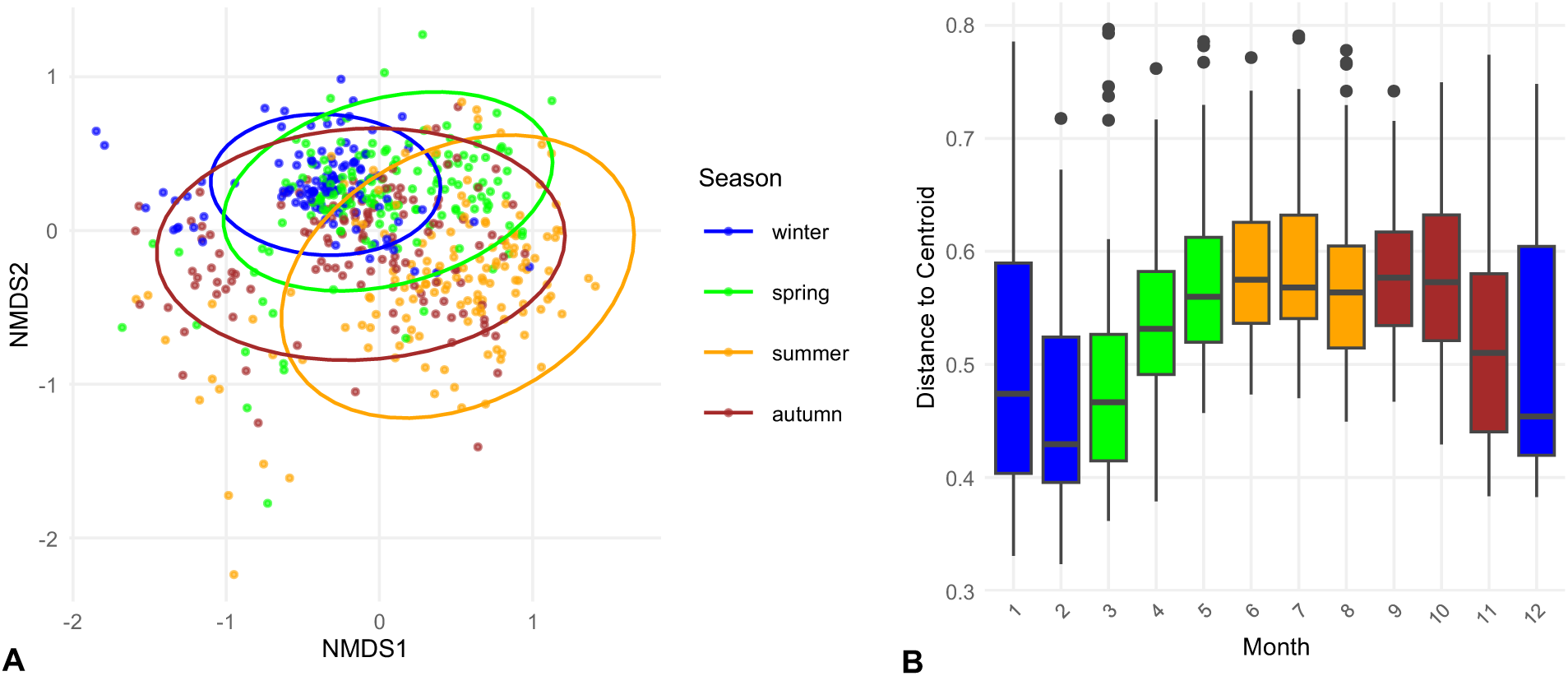
Intra-annual eukaryote diversity at Western Channel Observatory coastal site ‘L4’ across years (2001-2023; eDNA metabarcoding 18S-V9 rRNA), coloured by season (Winter: blue; Spring: green; Summer: yellow; Autumn: red): **A.** NMDS ordination (K = 3) of Bray-Curtis Dissimilarity (stress = 0.16; ADONIS test; ellipses 95% confidence interval). **B.** Dispersion plot to show the distance/dissimilarity of samples from their monthly ‘centroid’. Points indicate observations beyond 1.5 x the interquartile range.

Taxonomic classification was performed by DADA2 using the Protist Ribosomal Reference (PR^2^) database version 5.0.0 (Guillou *et al*., 2013). Eleven samples were sequenced twice to due to low success on the initial run; read counts were summed per sample. Taxonomy and abundance of ASVs were primarily interrogated using the R (R Core Team, 2004) package Phyloseq (McMurdie and Holmes, 2013). ASVs that were classified as bacteria or archaea (n = 1584) were removed from downstream analysis. DNA extraction and sequencing negative controls were examined to identify ASVs likely resulting from contamination using the R package Decontam (Davis *et al*, 2018). Of the 61 ASVs flagged, 52 were removed by subsequent pruning steps (i.e. removal of negative control/extraction kit blanks and very rare ASVs as described below). The remaining nine ASVs were further investigated via the NCBI nucleotide BLAST database (version: BLAST+ 2.16.0), plus evaluation of prevalence across samples and presence in low-abundance samples only. Three of the nine ASVs identified by Decontam were judged as not being contamination and were retained. After removal of negative controls (i.e. blank kit-and sequencing samples), very rare ASVs identified in <2 samples and <3 reads (n = 3443) were also removed (Caracciolo *et al*., 2022).

Further investigation of some abundant ASVs classified only as ‘Domain: Eukaryota’ was undertaken via NCBI BLAST, revealing that ASVs with high similarity (95%-98%) to archaeal sequences had been identified as eukaryotic by PR2. For conservatism, ASVs not characterised beyond domain level (n = 816) were removed. Per-sample read abundance was assessed and samples with <1000 reads (n = 41) were discarded (lowest Goods Coverage: 97.7%; remaining samples: n = 534). Due to inherent biases of PCR amplification and unavoidable differences in sequencing depth between samples, the number of sequences corresponding to the remaining 2980 ASVs were transformed to proportional values (i.e. relative abundance). Further, to facilitate downstream exploration of the potential influence of sequencing depth on the diversity patterns observed, a subsampled/rarefied version of the dataset was generated (at 1145 reads per sample, to match the lowest read number of the samples retained; Vegan, function: ‘rrarefy’) and transformed to relative abundance.

### Diversity indices and statistical tests

To explore seasonal variability of plankton alpha diversity, Shannon diversity, Simpson diversity and rarefied ASV richness were calculated per sample using the R package Vegan (Oksanen, 2001), and linear mixed-effects models (LMMs) were used to calculate the effect of season on alpha diversity metrics using the R package lme4 (Bates *et al*., 2015) (function: ‘lmer’; season: fixed effect; year: random intercept, to account for repeated measures and gaps). Because this was a combined time series, the potential confounding variables of sequencing run and sampling method deviations (i.e. in years following 2019 (38 samples): differences in minimum fraction size, collection method and volume filtered; across the time series: differences in filter material) were included as fixed effects in the models. Significance of seasonal variance was tested using Type III ANOVA with Satterthwaite’s method from lmerTest (Kuznetsova *et al*, 2017) (function: anova(m)). Seasonal estimated marginal means and post-hoc pairwise comparisons (Holm-adjusted) were conducted with R package emmeans (functions ‘emmeans()’ and ‘pairs()’, respectively) (Lenth and Piaskowski, 2025). Simpson diversity values underwent logit transformation to avoid violating assumptions of the model and ASV richness values were calculated from the subsampled/rarefied version of the dataset.

To investigate changes in plankton alpha diversity across the time series, adjusted annual means of plankton alpha diversity (Shannon; Simpson; rarefied AVS Richness) were computed after fitting LMMs (with season and potential confounding variables accounted for as fixed effects, and year as a random intercept), with year-specific random effect estimates extracted via ‘Best Linear Unbiased Predictions’ (BLUPs; function ‘ranef’). Year-level granularity was chosen (i.e. annual means, as opposed to monthly means or per-sample diversity) to facilitate the detection of inter-annual changepoints in this time series and to prevent the lack of samples for some year-months (Supporting Table 1) confounding the analysis. Changepoints in the adjusted annual mean series (mean/variance shifts) were calculated using the R package Changepoint (Killick and Eckley, 2014) (function: ‘cpt.meanvar’; method: ‘PELT’ (Pruned Exact Linear Time)).

Inspection of intra-annual (seasonal and monthly) plankton beta (community) diversity was undertaken using both relative abundance and presence-absence data, and rarefied versions of both data types. Bray-Curtis and Jaccard dissimilarity were calculated using the R package Vegan (function: ‘vegdist’; methods: “bray” and “jaccard” (respectively)), as were seasonal and monthly beta diversity dispersion analyses (functions: ‘betadisper’ and ‘permutest’). Vegan was also used for Nonmetric Multidimensional Scaling (NMDS; function: ‘metaMDS’). Differences in the dispersion of samples (i.e. distance between samples in Bray-Curtis dissimilarity) from the main periods of water column stratification and outside of this period were also tested with R package Vegan (functions: ‘betadisper’ and ‘permutest’). To control for the effect of potential confounding factors on plankton community diversity whilst disaggregating the effects of season and year, Distance-based Redundancy Analysis (db-RDA) models (package Vegan; function: ‘dbrda’) were generated on Bray-Curtis dissimilarity transformed relative abundance data. Significance was evaluated via permutation tests (999 permutations).

To examine the underlying structure of the plankton community data without *a priori* focus on the effect of season or year, Hierarchical Agglomerative Clustering (HAC) was performed on distance matrices of Bray-Curtis dissimilarity and – for further investigation of appropriate Average Silhouette Width (ASW) and visualisation - NMDS scores from Bray-Curtis dissimilarity and Jaccard dissimilarity (function: ‘hclust’; method: ‘ward.D2’). Average Silhouette Width (ASW) from R package library Cluster (Maechler *et al*, 2026) (function: ‘silhouette’) was used to assess the success of clustering when different solutions (i.e. 2-4 clusters) were applied (function: ‘cutree’). Exploration of potential abiotic factors effecting plankton diversity were conducted as follows: differences in the temperatures and NAO index values associated with HAC sample clusters were tested for significance using Kruskal-Wallis rank sum tests (function: ‘kruskal.test’). Correlations between alpha (Shannon) diversity and temperature/NAO index values were investigated with the function ‘cor.test’. Correlations between beta (community) diversity and temperature/NAO index values were tested using the vegan function ‘envfit’ (999 permutations) on a NMDS (k = 3, trymax = 100) of Bray-Curtis dissimilarity.

To examine plankton community cyclical dynamics in a multidimensional context, we conducted a Cyclical Ecological Trajectory Analysis (CETA) with the R package ecotraj (De Cáceres *et al*., 2019; Sturbois *et al*., 2021; Sturbois *et al*, 2023; Djeghri *et al*., 2026). CETA was performed on mean monthly distance scores from a Principal Coordinate Analysis (PCoA; R package: ape (Paradis and Schliep, 2019); function: ‘pcoa’) of Bray-Curtis dissimilarities - distance scores were agglomerated by month to capture major seasonal patterns and to minimise the effect of missing samples. Cycles containing ≥6 ecological states (i.e. months) were extracted (function: extractCycles; minEcolStates = 6) and were visualised via Principle Coordinates Analysis (PCoA). The first year (2001) was excluded from cycle extraction due to lack of sufficient boundary observations. Distances between cycles – a measure which accounts for cycle position, shape, size and direction - were calculated (function: ‘trajectoryDistances’, which utilises the distance metric Dsdsp), and the years driving the differences between the extracted cycles were evaluated (function: ‘dynamicVariation’).

Outputs from the analyses above were plotted using the R packages ggplot2 (Wickham, 2009), RColorBrewer (Neuwirth, 2022), patchwork (Pedersen, 2025) and ape (Paradis and Schliep, 2019). Data wrangling was assisted by R package tidyverse (Wickham *et al*, 2019).

## Results

### Seasonal succession and interannual shifts in plankton community structure

Over the two-decade time-series, plankton community composition exhibited clear temporal seasonal succession after accounting for interannual variation (dbRDA; adj. R² = 0.075, Pseudo-F = 16.9, df = 3, p = 0.001), with diversity higher in autumn and winter than in spring and summer (Supporting Table 2; Supporting Figures 2 and 3A). Controlling for potential confounding factors, including sequencing run and differences in sampling method, only slightly reduced the effect of season on community composition (dbRDA; adj. R² = 0.072, Pseudo-F = 16.2, df = 3, p = 0.001) (Supporting Figure 3B). Between years, communities were more similar during winter and early spring, whereas interannual variation was greater in summer and early autumn (Figure 1), and community composition also differed at monthly timescales (dispersion analysis of Bray–Curtis dissimilarity: seasons, F = 31.75, p = 0.001; months, F = 11.77, p = 0.001; Figure 1).

**Figure 2.**
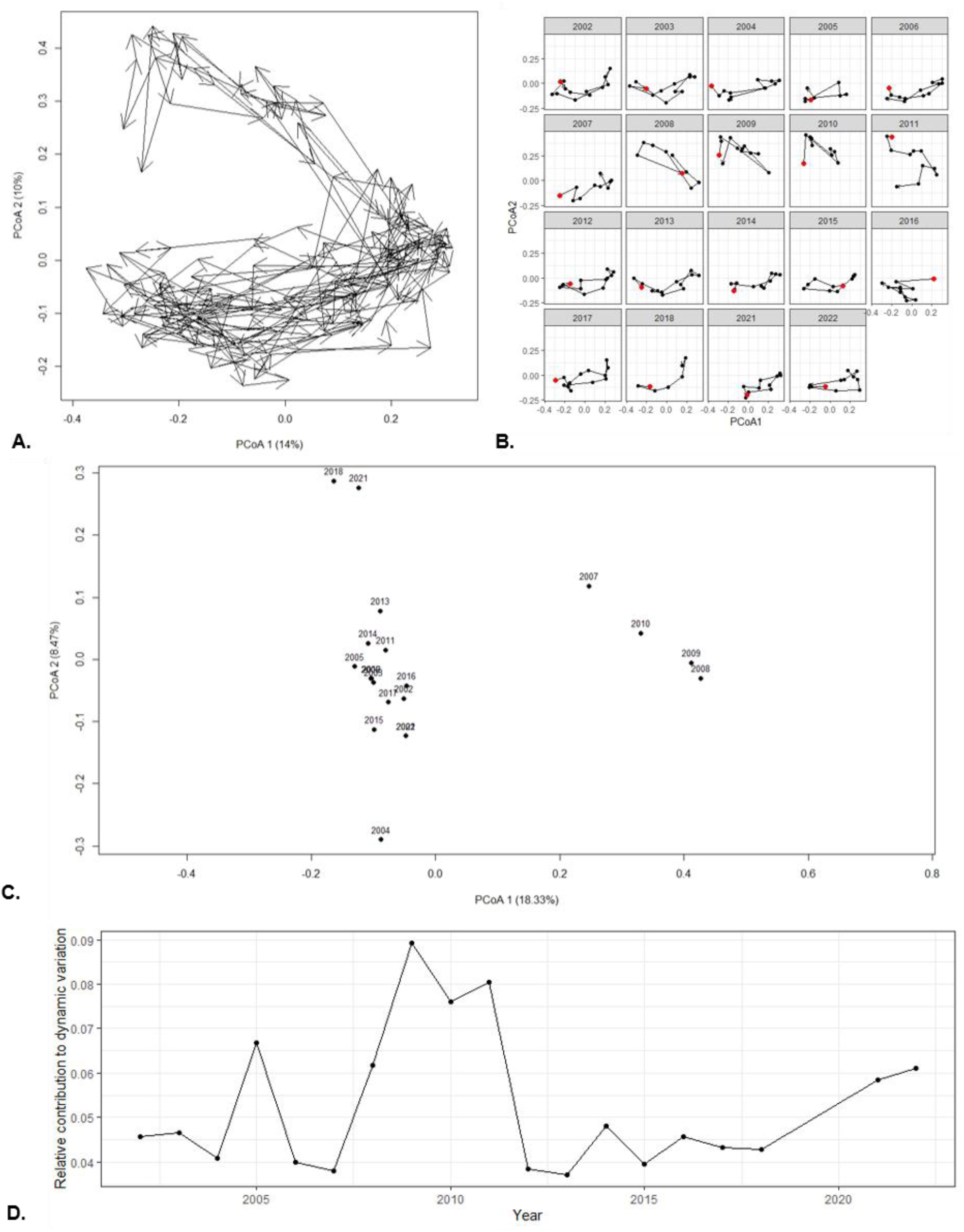
Cyclical Ecological Trajectory Analysis (CETA) of WCO-L4 plankton community (i.e. ASVs) time series: A. Ordination of all cycles in the timeseries; B. Cycles separated by year; initiation point coloured red; C. Visualisation (PCoA) of distances between cycles; points are labelled by cycle year; D. Relative contribution to cycle differences (i.e. dynamic variation) across years (2023 was excluded due to insufficient sample size for calculation of yearly cycle).

**Figure 3.**
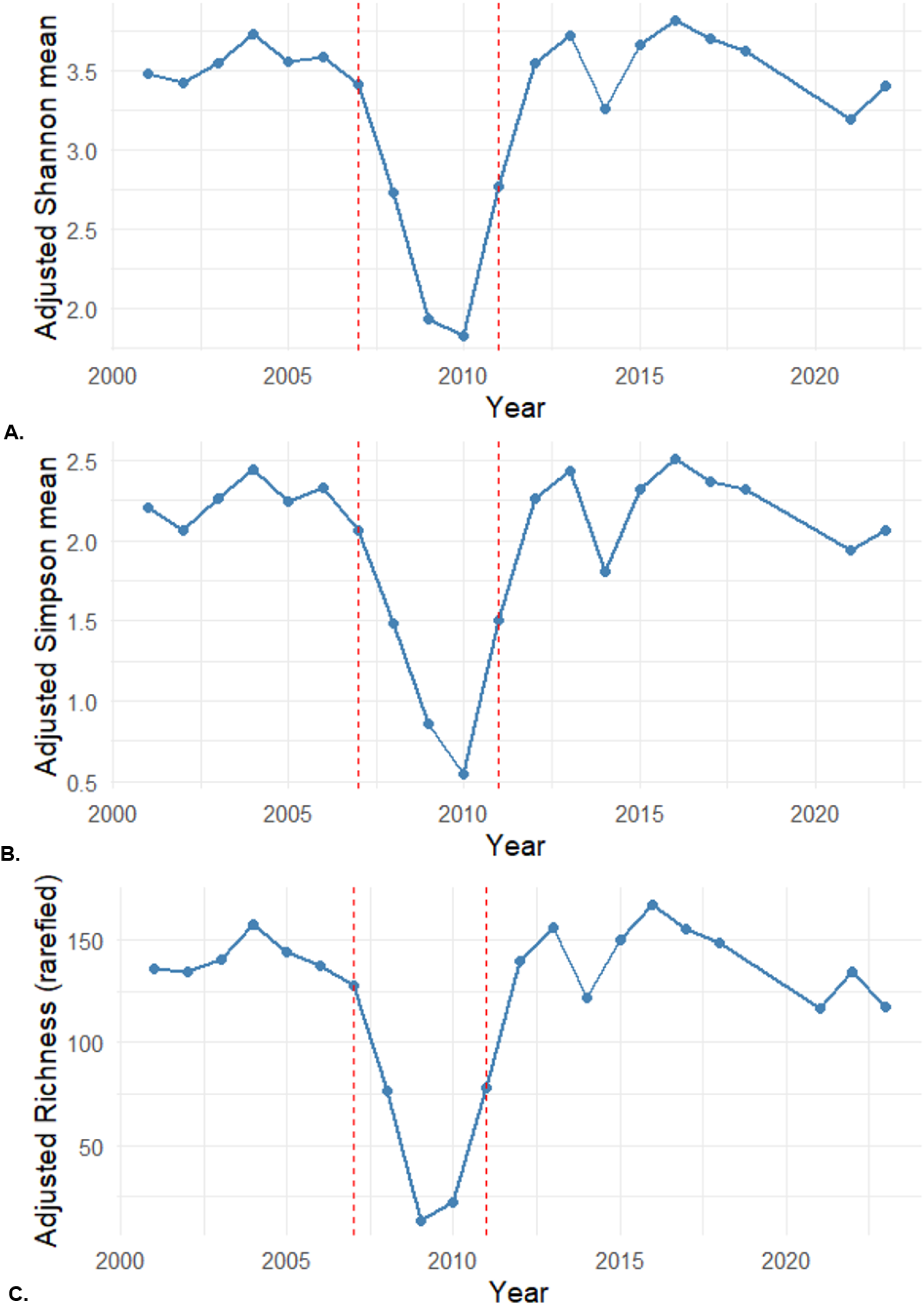
Inter-annual eukaryote diversity at Western Channel Observatory coastal site ‘L4’ across years (2001-2022; eDNA metabarcoding 18S-V9 rRNA): A. Adjusted annual means of plankton alpha diversity across the time series: (A) Shannon; (B) Simpson; (C) Richness; (annual means were estimated using a linear mixed-effect model with year included as a fixed factor and adjusted for season and sequencing run; 2023 was excluded due to insufficient sample size for calculation of annual mean. Dashed lines indicate regime ‘changepoints’ in the adjusted mean series (mean/variance shifts) using PELT).

**Figure 4.**
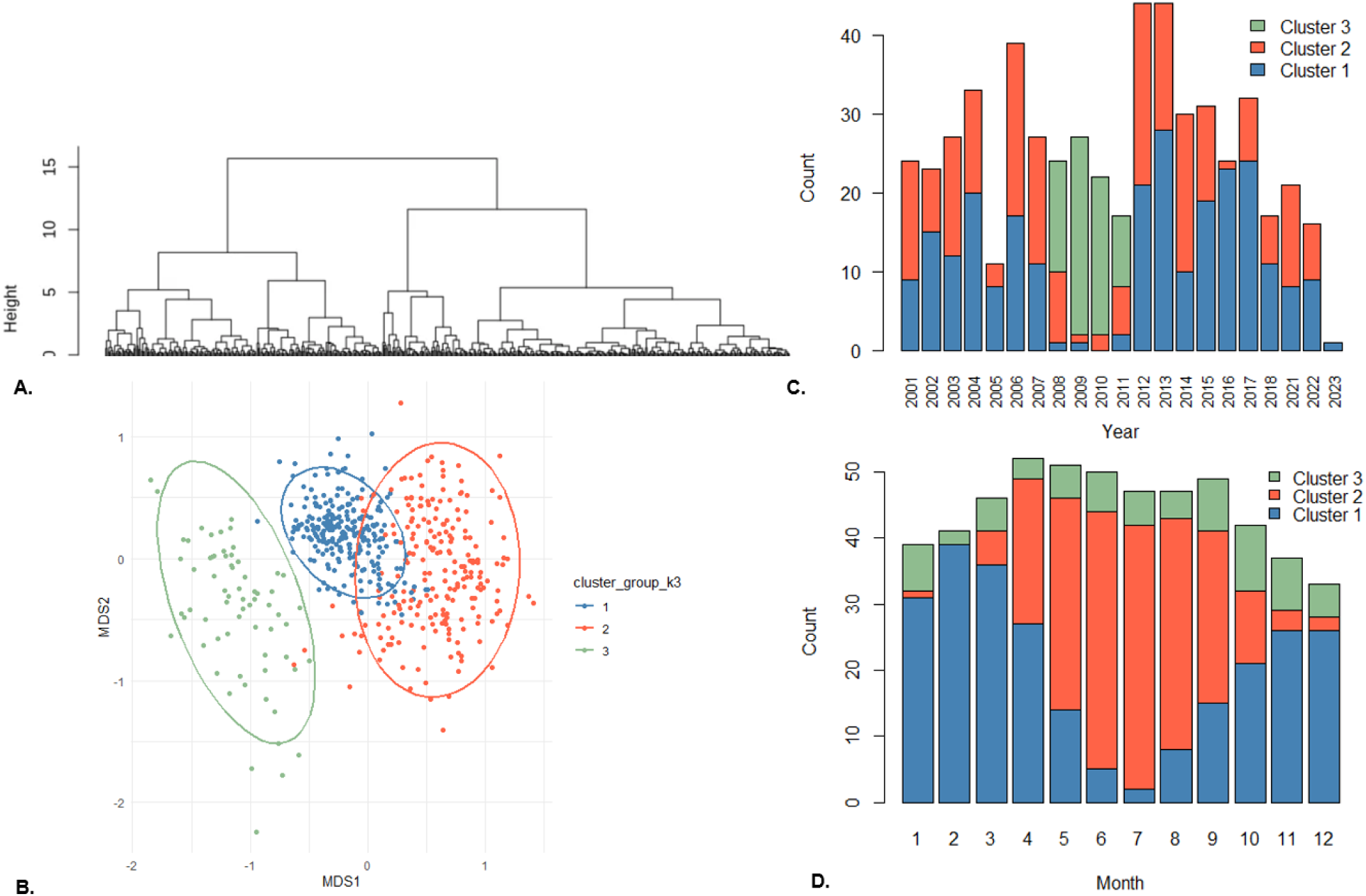
Inter-annual eukaryote diversity at Western Channel Observatory coastal site ‘L4’ across years (2001-2022; eDNA metabarcoding 18S-V9 rRNA) as revealed by Hierarchical Agglomerative Clustering of NMDS scores from Bray-Curtis dissimilarity; A. Cluster dendrogram; B. NMDS ordination of Bray-Curtis dissimilarity (K = 3); points are coloured to reflect hierarchical division into three clusters (ADONIS test; ellipses 95% confidence interval); Cluster 3 (coloured green; left of plot) is composed entirely of samples from the years 2008-2011 (Supporting Table 2); C. Sample counts per year, per hierarchical cluster; D. Sample counts per month, per hierarchical cluster.

The trajectories of seasonal cycles, determined using Cyclical Ecological Trajectory Analysis (CETA), overlapped for most years in the time series, however several years formed distinct trajectories that separated from the dominant cluster (Figure 2A, B). These shifted cycles occurred consecutively in the middle of the time series and included all seasons (Supporting Figure 5A, B). Analysis of distances among cycles showed that the years 2007–2010 were particularly dissimilar from the rest of the time series (Figure 2C), while dynamic variation among cycles was driven largely by the period 2008–2011, with a maximum in 2009 (Figure 2D). Together, these results indicate that over the 20-year time-series, substantial interannual variation in plankton community structure occurred, with communities from 2001 to ∼2007 appearing relatively similar, followed by a shift between ∼2008 and ∼2011, and then a return from ∼2012 onwards to a state more comparable to that observed before the shifted period.

**Figure 5.**
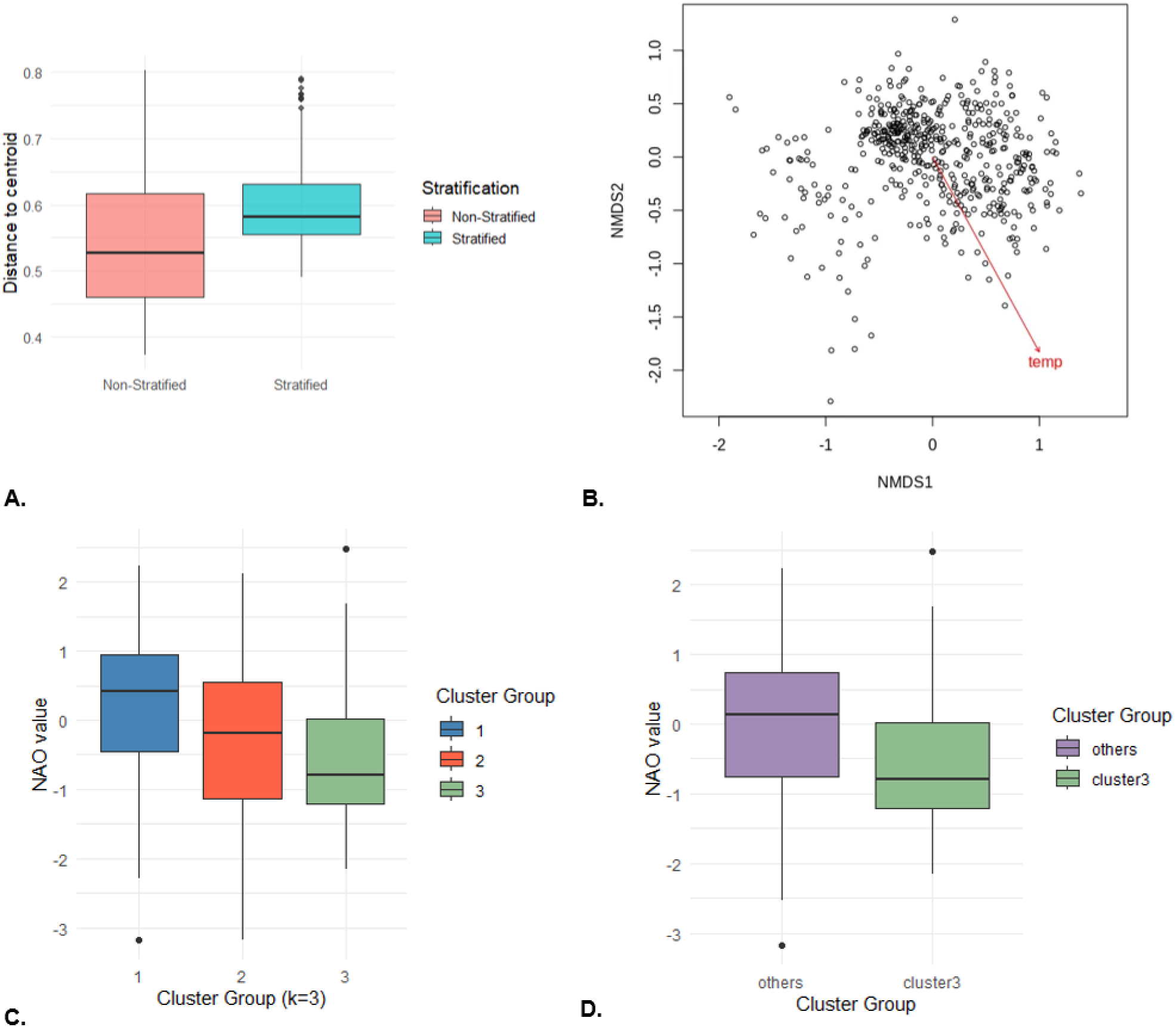
Candidate drivers of WCO-L4 plankton community diversity (total eukaryotes time series generated via metabarcoding; 18S-V9 rRNA marker); A. Dispersion plot to show the distance/dissimilarity of samples to the ‘centroid’ (i.e. the average of all samples) taken during the annual stratification period (May-September) and samples taken outside of this period. B. Envfit analysis on correlation of temperature with community (ASV) diversity (conducted on NMDS of Bray-Curtis dissimilarity). North Atlantic Oscillation (NAO) values across: C. Three groups of samples identified via Agglomerative Hierarchical Clustering; D. NAO values associated with ‘Cluster 3’ samples and all other samples.

### Interannual plankton biodiversity turnover

To investigate the apparent biodiversity anomaly in the years around 2010, we further analysed annual patterns in diversity and community composition. Alpha diversity, including Shannon diversity, Simpson diversity and ASV richness, was reduced during the years around 2010 (Supporting Figure 6A–F). A changepoint analysis (PELT) of adjusted annual means identified 2007 and 2011 as changepoints for all three alpha diversity metrics (Figure 3A-C). To quantify interannual variation in community composition, we conducted a second dbRDA analysis: after accounting for seasonal variation, year explained approximately 11% of the variation in plankton community composition (adj. R² = 0.111, F = 4.6, df = 20, p = 0.001). When potential confounding factors, including sequencing run and sampling method, were also controlled for, the effect of year remained significant, although the proportion of explained variance was slightly lower (adj. R² = 0.092, F = 4.2, df = 19, p = 0.001). In support of the early observations, the ordinations of both dbRDA models showed clear separation of the years 2008–2011 from the remainder of the time series (Supporting Figure 7A and B).

**Figure 6.**
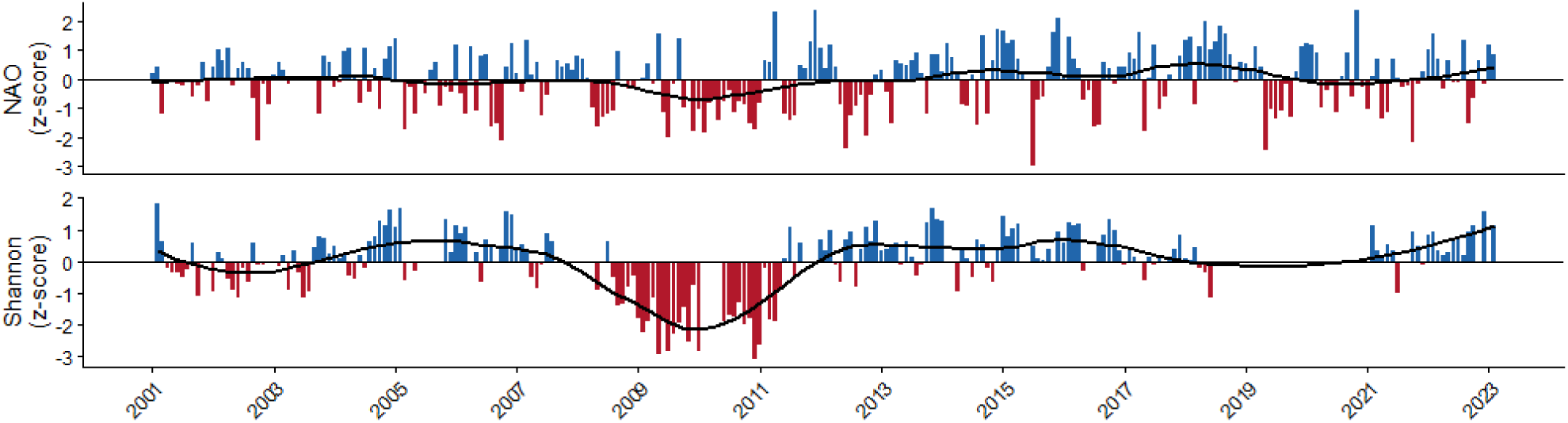
Monthly Z-scores with Locally Estimated Scatterplot Smoothing (LOESS) line for the (top) North Atlantic Oscillation (NAO) index and (bottom) Shannon Diversity at Western Channel Observatory coastal site ‘L4’ from a long-term (years 2001-2023) eDNA time series (metabarcoding for total eukaryotes;18S-V9 rRNA marker).

**Figure 7.**
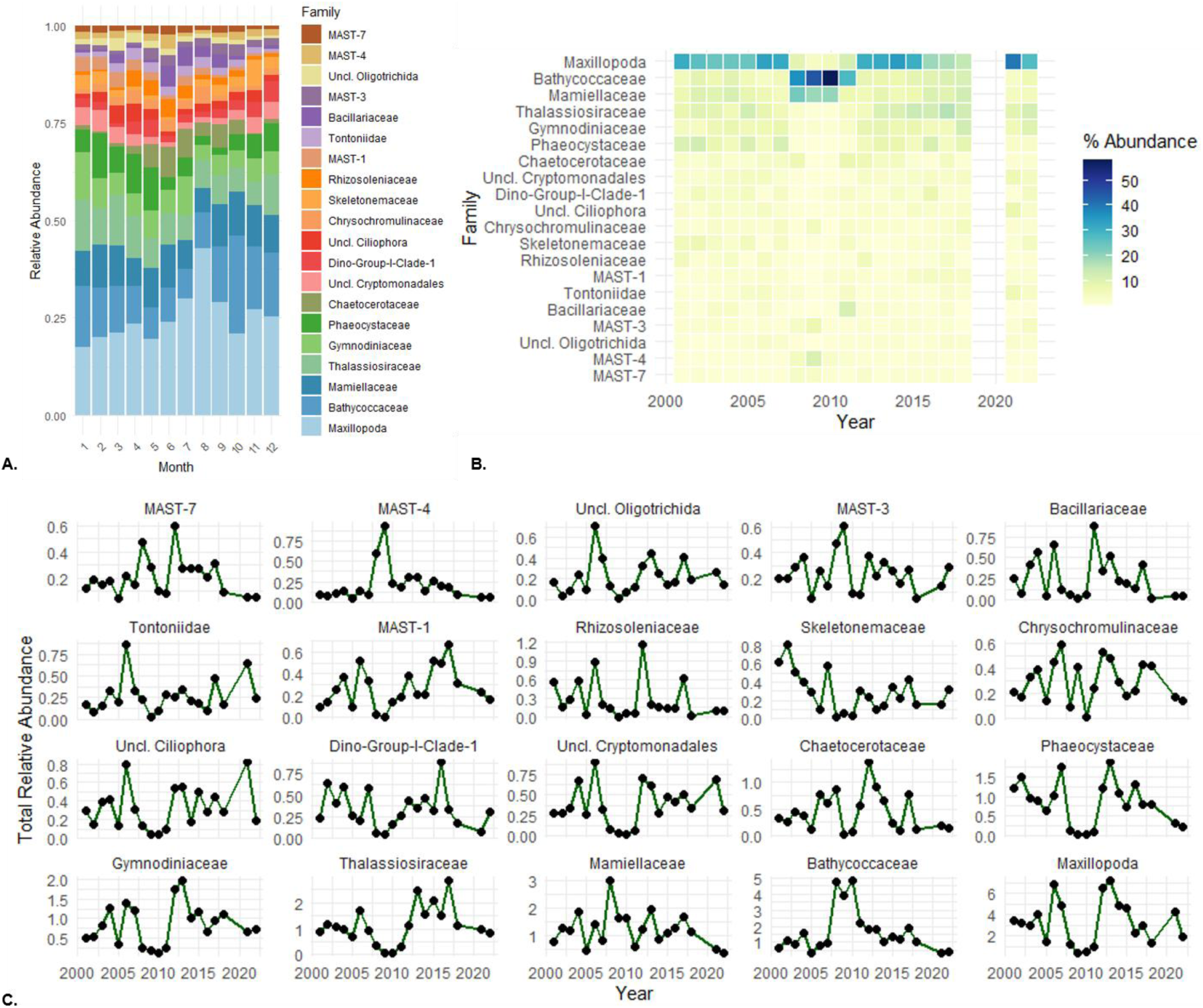
Relative abundance of the top twenty most abundant plankton families at Western Channel Observatory coastal site ‘L4’ from a long-term eDNA time series (metabarcoding for total eukaryotes;18S-V9 rRNA marker). A. Monthly relative abundance; B. Yearly mean relative abundances (years 2001-2022; 2023 was excluded due to insufficient sample size for calculation of yearly mean), normalised as percentage of total abundance of top twenty families (showing compositional shifts); C. Total mean relative abundance of each of the top twenty families over time (showing abundance trajectories).

To help identify specific timepoints/samples associated with this shifted period and further understand the underlying structure of the data, we applied Hierarchical Agglomerative Clustering (HAC) to distance matrices of Bray-Curtis dissimilarity. The resulting dendrogram indicated support for two to four clusters (Supporting Figure 8), however the highest Average Silhouette Width (ASW) obtained was surprisingly low (K = 2-4, ASW = 0.07-0.06). Application of HAC to sample scores from NMDS ordinations of Bray–Curtis dissimilarity also indicated support for two to four clusters (Figure 4A), with the highest (though still moderate) ASW obtained for three or four groups (K = 3-4, ASW = 0.31). The Bray-Curtis dissimilarity matrix was well preserved by the 3-dimentional NMDS ordination (Spearman’s ρ = 0.92), suggesting that inaccurate representation of the data by the NMDS was not an issue. The low ASW values arising from the clustering of Bray-Curtis dissimilarities may result from both the high degree of dimensionality in the data, and the narrow, high range of within-cluster and between-cluster Bray-Curtis distances (0.69-0.94; Supporting Table 3). In support of genuine clusters within the gradient of variation observed (Figure 4B), between-cluster distances exceeded within-cluster distances for every pairwise comparison (Supporting Table 3).

In the three-cluster solution (in both HAC analyses), Cluster 3 (Figure 4B) consisted exclusively of samples from 2008–2011 (Figure 4C; Supporting Table 2A) and included samples from across the year (Figure 4D; Supporting Table 2B). By contrast, Clusters 1 and 2 differed strongly in the proportions of samples associated with the main stratified period versus the remainder of the year (Figure 4D; Supporting Table 2B). In comparison to Cluster 2 - comprised largely of samples from the main water column stratified period (i.e. May – September), Cluster 1, which had a high proportion of samples from October to April (Figure 4D; Supporting Table 2B), included very few samples taken from 2008-2011 (Figure 4C; Supporting Table 2A), suggesting that these periods were of particular importance to the formation of cluster 3.

A further HAC analysis of sample scores from an NMDS ordination of Jaccard dissimilarity also produced a separate cluster comprised of samples from 2008-2011 (2-4 cluster solutions; K = 2, ASW = 0.51; K = 3, ASW = 0.36; K = 4, ASW = 0.30; Supporting figure 9), indicating that the ‘shift’ period in plankton community composition is not only due to differences in taxa abundance, but also due to differences in taxa presence-absence/turnover. Patterns observed in HAC analyses were retained following rarefaction to a common sequencing depth (i.e. subsampling of sequences) (Supporting Figures 10 and 11), indicating that differences in sample coverage were not responsible for the patterns observed.

### Local hydrography and basin-scale climate forcing as candidate drivers

Water-column structure appeared to be an important factor for plankton community composition in the time series (Figure 5A; Supporting Table 4B). This pattern is consistent with the recurring stratification cycle at Station L4 and the associated pelagic dynamics. Across the time series, plankton communities were more dissimilar under stratified than mixed water-column conditions (F = 62.55, p < 0.001; Figure 5A). Temperature was also associated with community composition, with envfit analysis on the NMDS ordination of Bray–Curtis dissimilarity revealing a correlation (R² = 0.397, p = 0.001; Figure 5B). The fitted temperature vector aligned closely with the seasonal distribution of samples in ordination space, indicating that temperature is an important driver of the observed seasonal community structure (Figure 1A; Figure 5B).

Temperature also differed among the three sample clusters identified by hierarchical clustering (Kruskal–Wallis χ² = 100.08, df = 2, p < 0.001; Supporting Figure 12A). However, temperature alone does not appear to explain the anomalous proposed biodiversity turnover period. In particular, temperatures associated with Cluster 3 samples did not differ from those of other samples (Kruskal–Wallis χ² = 0.05, df = 1, p = 0.824; Supporting Figure 12B), and the correlation between temperature and Shannon diversity was weak (Pearson’s r(500) = −0.118, p < 0.01, CI [−0.203, −0.030]; Supporting Figure 12C). However, the NAO was more closely associated with the anomalous sample group. NAO index values differed among the three sample clusters identified by hierarchical clustering (Kruskal–Wallis χ² = 41.14, df = 2, p < 0.001; Figure 5C), and Cluster 3 samples were associated with lower, more negative NAO values than the other samples (Kruskal– Wallis χ² = 11.853, df = 1, p < 0.001; Figure 5D). Consistent with this pattern, Shannon diversity showed a weak but significant positive correlation with NAO index values (Pearson’s r(532) = 0.137, p < 0.01, CI [0.053, 0.219]; Supporting Figure 13), and both NAO and Shannon diversity values underwent a sustained negative period around 2010 (Figure 6). Together, these results suggest that while seasonal-scale community structure is closely linked to local hydrographic conditions (i.e. stratification and seawater temperature), the transient biodiversity turnover observed during 2008– 2011 is more consistent with basin-scale climate forcing associated with negative NAO phases.

### Major plankton families and the biodiversity turnover

Over the time series, the top twenty most abundant plankton families (Figure 7A) included copepods, diatoms, dinoflagellates, pico-algae and ciliates. These observations are congruous with previous reports of diatoms, dinoflagellates, larger nanoflagellates (≥15 µm) and copepods as dominating Station L4 biomass datasets enumerated via light microscopy and flow cytometry (McEvoy *et al*, 2023). The long-term patterns of relative abundance of the top twenty most abundant plankton families in the Station L4 eukaryote time series show taxa specific responses across the years of anomalous diversity (Figure 7B and C). Groups that underwent an apparent reduction in relative abundance during the 2008-2011 period include some of the relatively larger taxa, including zooplankton (Maxillopoda: crustaceans; largely copepods), and some larger phytoplankton (e.g. Thalassiosiraceae: diatoms; Gymnodiniaceae: dinoflagellates). Conversely, the relative abundance of some picoplankton groups peaked in the anomalous years, including phytoplankton (e.g. Bathycoccaceae and Mamiellaceae: green algae) and heterotrophs (MAST-3; MAST-4; MArine STramenopiles).

## Discussion

This study uses a two-decade 18S rRNA gene (V9) metabarcoding time series from a temperate coastal observatory to resolve eukaryotic plankton dynamics across seasonal to interannual timescales. Plankton community composition showed a recurrent seasonal structure, consistent with the well-established annual cycle of the Western English Channel and with previous molecular and observational studies at Station L4 and comparable coastal systems (Gilbert *et al*., 2009; Gilbert *et al*., 2012; Lambert *et al*., 2019; McEvoy *et al*., 2023). Superimposed on this seasonal template was a distinct period of biodiversity turnover during approximately 2008–2011, characterised by reduced alpha diversity and altered community composition. This reorganisation was transient, with communities after 2011 returning towards a state more comparable with the earlier part of the record. Together, these findings indicate that long-term metabarcoding can reveal meaningful reorganisations that may be obscured in shorter studies or in monitoring focused on biomass or larger plankton alone.

Seasonal succession was the clearest and most persistent structure in the eukaryotic plankton community. This is expected in a temperate shelf-sea system in which light, temperature, nutrient availability and water-column stability vary strongly through the year. In the Western English Channel, seasonal stratification typically develops from late spring through early autumn, producing temporal windows in which resource supply, grazing pressure and dispersal differ from mixed winter conditions (McEvoy *et al*., 2023). The stronger interannual similarity of winter and early-spring communities, relative to summer and autumn communities, suggests that mixed conditions may repeatedly reset parts of the plankton assemblage, while stratification permits more variable successional pathways. This interpretation is consistent with previous work showing recurrent seasonality in plankton communities, both at L4 and elsewhere, despite substantial short-term environmental variability (Gilbert *et al*., 2009; Gilbert *et al*., 2012; Lambert *et al*., 2019).

A major result is the detection of a discrete biodiversity anomaly centred on 2008–2011. Multiple lines of analysis support this interpretation including changepoints in adjusted annual diversity metrics, separation of these years in dbRDA ordinations, distinct seasonal cycles in CETA, and the recovery of a 2008–2011 sample group by hierarchical clustering. Importantly, the same period was apparent in both Bray–Curtis and Jaccard analyses, indicating that the anomaly was not solely a redistribution of relative abundances among persistent taxa but also involved turnover in taxon occurrence. The convergence among methods strengthens the inference that this was a community-wide reorganisation rather than an artefact of a single analytical approach. However, the return of post-2011 communities towards the pre-shift domain suggest that the system did not undergo a persistent regime shift in the strict sense. Instead, the data are more consistent with transient biodiversity turnover (Pinsky *et al*., 2025). This finding aligns with evidence that biodiversity change is often expressed as temporal turnover. Recent syntheses have highlighted that warming and cooling can both catalyse compositional change, with communities reorganising as taxa track shifting environmental conditions (Pinsky *et al*., 2025).

The association between the shifted sample group and negative NAO values suggests that basin-scale atmospheric forcing contributed to the 2008–2011 anomaly. The NAO influences North Atlantic and European shelf ecosystems through effects on wind stress, winter temperature, storm frequency, vertical mixing and nutrient supply (Fromentin and Planque, 1996; Hurrell *et al*., 2003; Gordo *et al*., 2011). At L4, temperature was strongly aligned with seasonal community structure but did not distinguish the anomalous cluster from the remainder of the time series. By contrast, NAO values were lower for the shifted samples, and winter samples contributed strongly to the distinctiveness of the period. This points towards an indirect climate-forcing mechanism in which altered winter conditions changed the physical and nutrient context from which subsequent seasonal succession developed.

Previous studies support such a link between NAO variability and plankton composition. In the English Channel, spring bloom composition has been related to the NAO, with higher diatom contribution under positive NAO conditions (Irigoien *et al*., 2000). Across the North Atlantic, NAO variability has also been associated with shifts in phytoplankton size structure and organic carbon flux (Henson *et al*., 2012), as well as changes in copepod distributions and abundance (Fromentin and Planque, 1996; Piontkovski *et al*., 2006). The L4 metabarcoding record extends these observations by showing that a negative NAO interval was associated with whole-community eukaryotic turnover, encompassing phytoplankton, heterotrophic protists and metazoan taxa. This broader community perspective is important because changes in one plankton component can propagate through grazing, competition, parasitism and trophic coupling (Kasada *et al*., 2026).

The inferred shift towards smaller plankton and microbial food-web components has potential consequences for ecosystem function. Larger diatoms and copepods contribute substantially to food-web transfer and carbon export (Tréguer *et al*., 2018), whereas smaller picoeukaryotes and heterotrophic protists are more tightly embedded in microbial recycling pathways (Worden *et al*., 2015). A relative increase in small cells may reduce the efficiency with which primary production is transferred to larger grazers or exported as sinking particles, although the outcome depends on aggregation, grazing interactions and trophic routing (Moriceau, *et al,* 2018; Tréguer *et al*., 2018). Because plankton underpin marine food webs and biogeochemical cycling, transient community reorganisation can have ecological consequences even when it does not persist as a long-term regime shift (Grigoratou et al., 2025; Siegel et al., 2026).

At the same time, caution is required when interpreting marker-gene relative abundance. The 18S-V9 marker provides broad taxonomic coverage but is affected by variation in rRNA gene copy number, cell size, primer affinity and DNA persistence (Zhu *et al*., 2005). As a result, relative read abundance is not equivalent to cell abundance or biomass. The strength of the present study lies instead in its capacity to detect coherent temporal structure across many taxa and samples. Future integration with microscopy, flow cytometry, pigment data, zooplankton counts and biogeochemical measurements from the Western Channel Observatory would allow the functional consequences of the 2008–2011 turnover to be tested more directly (Tarran and Bruun, 2015; McEvoy *et al*., 2023).

In conclusion, this metabarcoding time series demonstrates that coastal eukaryotic plankton communities are structured by strong recurrent seasonality but can also undergo marked, multi-year biodiversity turnover associated with basin-scale climate forcing. The 2008–2011 anomaly was characterised by reduced diversity, altered community composition and a relative shift from larger plankton groups towards smaller picoeukaryotic and heterotrophic components. Because the community subsequently returned towards its earlier state, the event is best interpreted as a transient reorganisation rather than a persistent regime shift. This distinction matters for understanding resilience. Coastal plankton communities may be highly responsive to climate variability yet retain the capacity for recovery when forcing changes. More broadly, the study highlights the importance of long-term molecular observatories for detecting episodic biodiversity change and for linking plankton community dynamics to climate variability, food-web structure and ecosystem function.

## Supporting information

Supporting Documentation

## Data Accessibility

The sequencing data generated in this study has been deposited in the European Nucleotide Archive (ENA) under the Study accession number PRJEB98106 (secondary accession number ERP180536).

## Author Contributions

Joanna Warwick-Dugdale performed the research, analysed the data, prepared figures and tables, authored and reviewed drafts of the paper and approved the final draft. Clare Ostle authored or reviewed drafts of the paper and approved the final draft. Kimberley Bird, Yousef Touhami, Rohan Allen and Cordelia Roberts performed the research, authored or reviewed drafts of the paper and approved the final draft. Laura Branscombe performed the research. Katherine Helliwell, Nicolas Henry and Nicolas Djeghri authored or reviewed drafts of the paper and approved the final draft. Tim Smyth contributed materials. Michael Cunliffe conceived and designed the research, contributed reagents/materials/analysis tools, authored and reviewed drafts of the paper and approved the final draft.

## Acknowledgments

The authors thank everyone who has contributed to the Western Channel Observatory, in particular the crews of the Plymouth Marine Laboratory RV ‘Quest’ and the Marine Biological Association RV ‘Sepia’ for the collection of water samples, and Professor Andy Rees for Station L4 nutrients data. Parts of the analyses undertaken here were conducted with the high-power computing resources provided by the Analysis and Bioinformatics for Marine Science (ABiMS) platform at the Roscoff Biological Station.

