## Supporting Documentation for "Biodiversity turnover in coastal marine eukaryotic plankton revealed by a 20-year metabarcoding time series"

**Supporting Table 1.** Number of samples per month and year across the L4 eDNA time series.

| Month/ Year | 1 | 2 | 3 | 4 | 5 | 6 | 7 | 8 | 9 | 10 | 11 | 12 | Total |
|---|---|---|---|---|---|---|---|---|---|---|---|---|---|
| **2001** | 0 | 1 | 1 | 2 | 4 | 3 | 3 | 1 | 3 | 2 | 4 | 0 | 24 |
| **2002** | 2 | 1 | 2 | 3 | 2 | 2 | 3 | 2 | 2 | 2 | 2 | 0 | 23 |
| **2003** | 0 | 1 | 1 | 3 | 3 | 3 | 3 | 4 | 4 | 2 | 2 | 1 | 27 |
| **2004** | 1 | 3 | 3 | 1 | 3 | 3 | 4 | 4 | 3 | 2 | 4 | 2 | 33 |
| **2005** | 2 | 1 | 1 | 2 | 1 | 0 | 0 | 0 | 0 | 0 | 2 | 2 | 11 |
| **2006** | 4 | 4 | 4 | 4 | 3 | 4 | 3 | 4 | 3 | 3 | 1 | 2 | 39 |
| **2007** | 4 | 3 | 4 | 4 | 2 | 3 | 4 | 3 | 0 | 0 | 0 | 0 | 27 |
| **2008** | 0 | 0 | 0 | 0 | 4 | 5 | 1 | 1 | 5 | 3 | 3 | 2 | 24 |
| **2009** | 4 | 1 | 2 | 1 | 1 | 5 | 3 | 2 | 2 | 3 | 1 | 2 | 27 |
| **2010** | 2 | 0 | 0 | 0 | 0 | 2 | 2 | 2 | 3 | 4 | 4 | 3 | 22 |
| **2011** | 1 | 1 | 3 | 3 | 1 | 3 | 1 | 1 | 3 | 0 | 0 | 0 | 17 |
| **2012** | 3 | 5 | 3 | 4 | 5 | 3 | 3 | 4 | 4 | 4 | 3 | 3 | 44 |
| **2013** | 3 | 4 | 5 | 4 | 3 | 4 | 5 | 4 | 3 | 3 | 3 | 3 | 44 |
| **2014** | 1 | 0 | 0 | 1 | 4 | 4 | 5 | 3 | 3 | 4 | 2 | 3 | 30 |
| **2015** | 3 | 3 | 3 | 4 | 0 | 0 | 3 | 4 | 5 | 4 | 0 | 2 | 31 |
| **2016** | 2 | 4 | 1 | 4 | 3 | 0 | 0 | 1 | 2 | 2 | 2 | 3 | 24 |
| **2017** | 2 | 2 | 4 | 3 | 4 | 1 | 1 | 5 | 3 | 2 | 2 | 3 | 32 |
| **2018** | 2 | 2 | 3 | 4 | 4 | 2 | 0 | 0 | 0 | 0 | 0 | 0 | 17 |
| **2021** | 0 | 2 | 4 | 4 | 2 | 2 | 2 | 1 | 0 | 1 | 2 | 1 | 21 |
| **2022** | 3 | 2 | 2 | 1 | 2 | 1 | 1 | 1 | 1 | 1 | 0 | 1 | 16 |
| **2023** | 0 | 1 | 0 | 0 | 0 | 0 | 0 | 0 | 0 | 0 | 0 | 0 | 1 |

**Supporting Table 2.** Results of Type III Analysis of Variance (ANOVA; Satterthwaite's method) and Post-hoc pairwise comparisons (Holm-adjusted) of Linear mixed-effect models (LMMs) on the effect of season (fixed effect) on three diversity indices: Shannon; Simpson; ASV Richness (rarefied/subsampled data), after accounting for year (as a random effect) and the potential confounding effect of sequencing run ('seq. run'; fixed effect), for coastal station 'L4' eDNA eukaryote (metabarcoding) time series. Global test model F and P values displayed; Different letters indicate significant differences in estimated marginal means.

| Metric | Season effect (F) | P (season) | Seq. run effect (F) | P (seq. run) | Seasonal ranking (estimated marginal means with significance groups) |
|---|---|---|---|---|---|
| Shannon | 11.05 | <0.001 | 3.16 | <0.01 | Winter (3.97) b ≈ Autumn (3.92) b > Summer (3.69) a ≈ Spring (3.65) a |
| Simpson* | 7.46 | <0.001 | 1.28 | 0.27 | Winter (2.94) b ≈ Autumn (2.92) b > Summer (2.63) a ≈ Spring (2.66) a |
| Richness# | 21.58 | <0.001 | 15.72 | <0.001 | Winter (169) c > Autumn (155) b > Summer (141) a ≈ Spring (133) a |

*Simpson values logit transformed; #Richness values from subsampled/rarefied dataset

**Supporting Table 3.** Within-cluster and between-cluster distances from hierarchical (agglomerative) clustering analysis on Bray-Curtis dissimilarities (K = 3).

| | within cluster | vs cluster 1 | vs cluster 2 | vs cluster 3 |
|---|---|---|---|---|
| Cluster 1 | 0.694 | — | 0.828 | 0.880 |
| Cluster 2 | 0.815 | 0.828 | — | 0.939 |
| Cluster 3 | 0.819 | 0.880 | 0.939 | — |

**Supporting Table 4.** Samples split according to a hierarchical (agglomerative) clustering analysis on per-sample community (ASV) diversity (i.e. NMDS scores of Bray-Curtis dissimilarity); **A).** Number of samples per year in each of three clusters; Cluster 3 is comprised of samples from 2008-2011 only (bold); **B).** Number of samples per month in each of three clusters; Cluster 2 includes a high proportion of samples from months when the water column is stratified (bold); Cluster 1 includes a high proportion of samples from the months when the water column is mixed (bold).

**A.**

| Year | Bray-Curtis Cluster 1 | Bray-Curtis Cluster 2 | Bray-Curtis Cluster 3 | NMDS Cluster 1 | NMDS Cluster 2 | NMDS Cluster 3 |
|---|---|---|---|---|---|---|
| 2001 | 06 | 18 | 0 | 09 | 15 | 0 |
| 2002 | 09 | 14 | 0 | 15 | 08 | 0 |
| 2003 | 10 | 17 | 0 | 12 | 15 | 0 |
| 2004 | 15 | 18 | 0 | 20 | 13 | 0 |
| 2005 | 06 | 5 | 0 | 08 | 03 | 0 |
| 2006 | 15 | 24 | 0 | 17 | 22 | 0 |
| 2007 | 03 | 24 | 0 | 11 | 16 | 0 |
| **2008** | 01 | 08 | **15** | 01 | 09 | **14** |
| **2009** | 01 | 01 | **25** | 01 | 01 | **25** |
| **2010** | 0 | 0 | **22** | 0 | 02 | **20** |
| **2011** | 02 | 04 | **11** | 02 | 06 | **09** |
| 2012 | 17 | 27 | 0 | 21 | 23 | 0 |
| 2013 | 18 | 26 | 0 | 28 | 16 | 0 |
| 2014 | 08 | 22 | 0 | 10 | 20 | 0 |
| 2015 | 14 | 17 | 0 | 19 | 12 | 0 |
| 2016 | 17 | 07 | 0 | 23 | 01 | 0 |
| 2017 | 17 | 15 | 0 | 24 | 08 | 0 |
| 2018 | 04 | 13 | 0 | 11 | 06 | 0 |
| 2021 | 07 | 14 | 0 | 08 | 13 | 0 |
| 2022 | 07 | 09 | 0 | 09 | 07 | 0 |
| 2023 | 01 | 0 | 0 | 01 | 0 | 0 |

**B.**

| Month | Bray-Curtis Cluster 1 | Bray-Curtis Cluster 2 | Bray-Curtis Cluster 3 | NMDS Cluster1 | NMDS Cluster2 | NMDS Cluster3 |
|---|---|---|---|---|---|---|
| 1 | **29** | 03 | 07 | **32** | 01 | 06 |
| 2 | **33** | 06 | 02 | **39** | 0 | 02 |
| 3 | **16** | 25 | 05 | **36** | 05 | 05 |
| 4 | 09 | 40 | 03 | **27** | 22 | 03 |
| **5** | 06 | **40** | 05 | 14 | **32** | 05 |
| **6** | 01 | **41** | 08 | 05 | **39** | 06 |
| **7** | 01 | **41** | 05 | 02 | **40** | 05 |
| **8** | 02 | **40** | 05 | 08 | **35** | 04 |
| **9** | 14 | **25** | 10 | 15 | **26** | 08 |
| 10 | **18** | 14 | 10 | **21** | 11 | 10 |
| 11 | **28** | 03 | 08 | **26** | 03 | 08 |
| 12 | **23** | 05 | 05 | **26** | 02 | 05 |

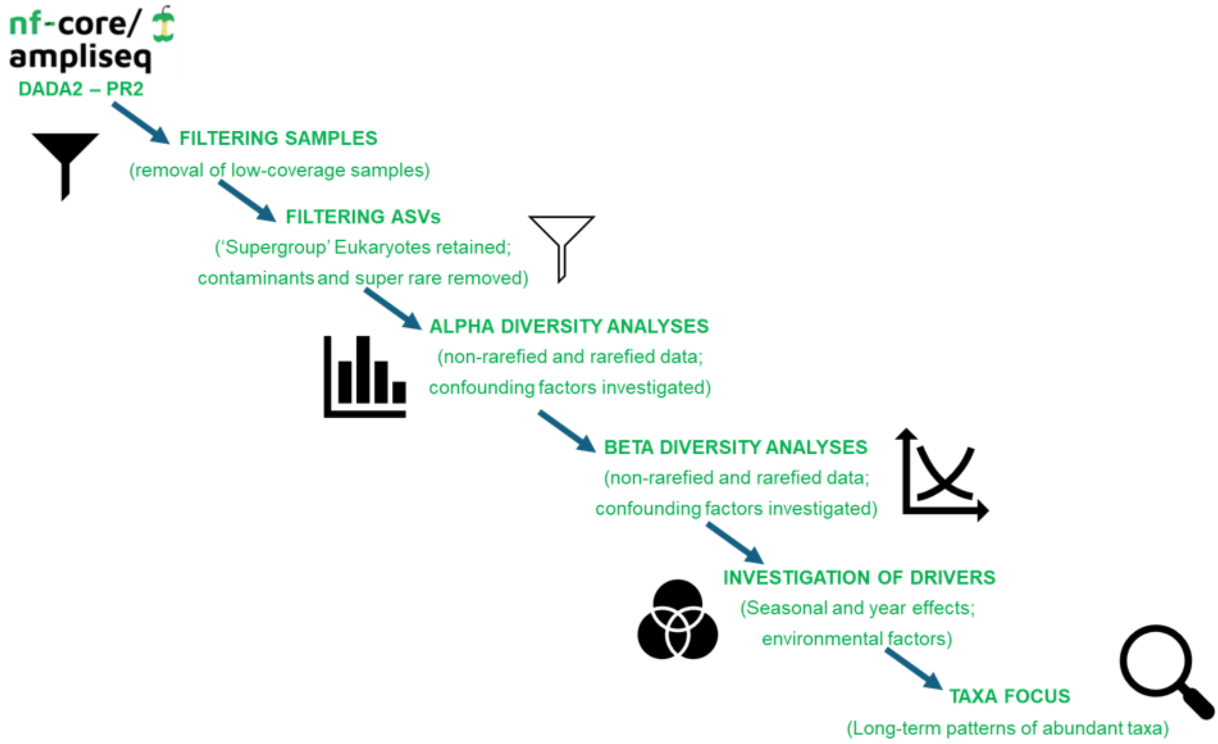


**Supporting Figure 1.** Summary of the bioinformatic pipeline employed to investigate the intra- and interannual patterns plankton biodiversity in the WCO-L4 metabarcoding (18S-V9 marker) time series.

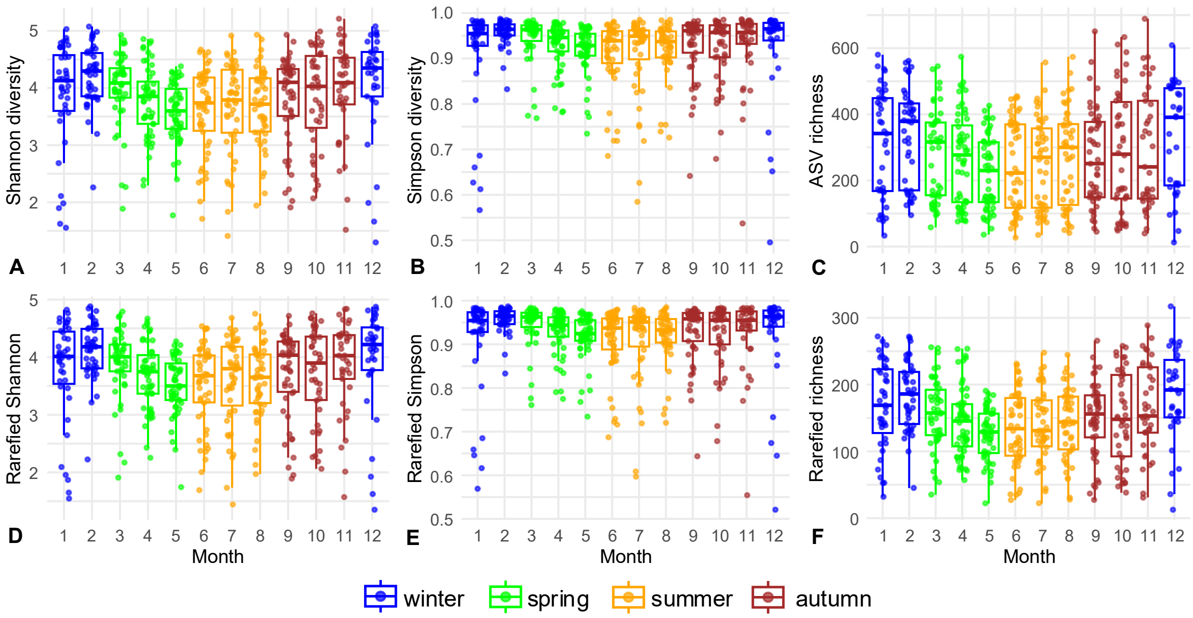


**Supporting Figure 2**. Intra-annual diversity: Monthly alpha diversity at Western Channel Observatory coastal site 'L4' from a long-term (years 2001-2023) eukaryote eDNA (metabarcoding: 18S-V9 rRNA) time series as calculated from the full dataset (A. Shannon; B. Simpson; C. ASV richness) and a subsampled/rarefied dataset (D. rarefied Shannon; E. rarefied Simpson; F.  rarefied ASV richness); coloured by season (Winter: blue; Spring: green; Summer: yellow; Autumn: red).

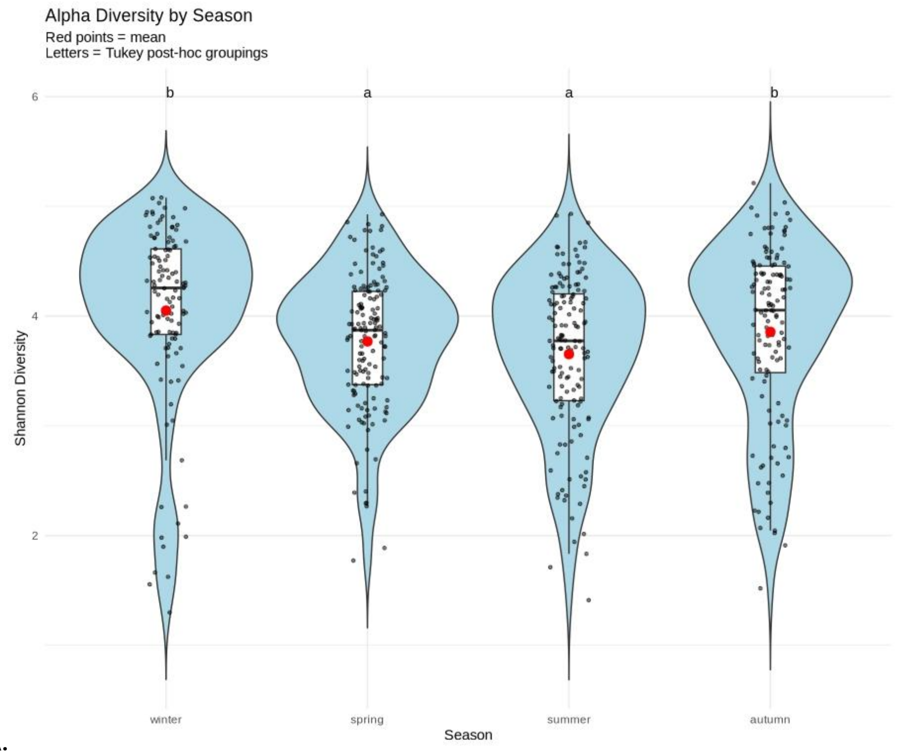
Alpha Diversity by Season
Red points = mean
Letters = Tukey post-hoc groupings
b
a
a
b
6
4
2
Shannon Diversity
winter
spring
summer
autumn
Season

**A.**

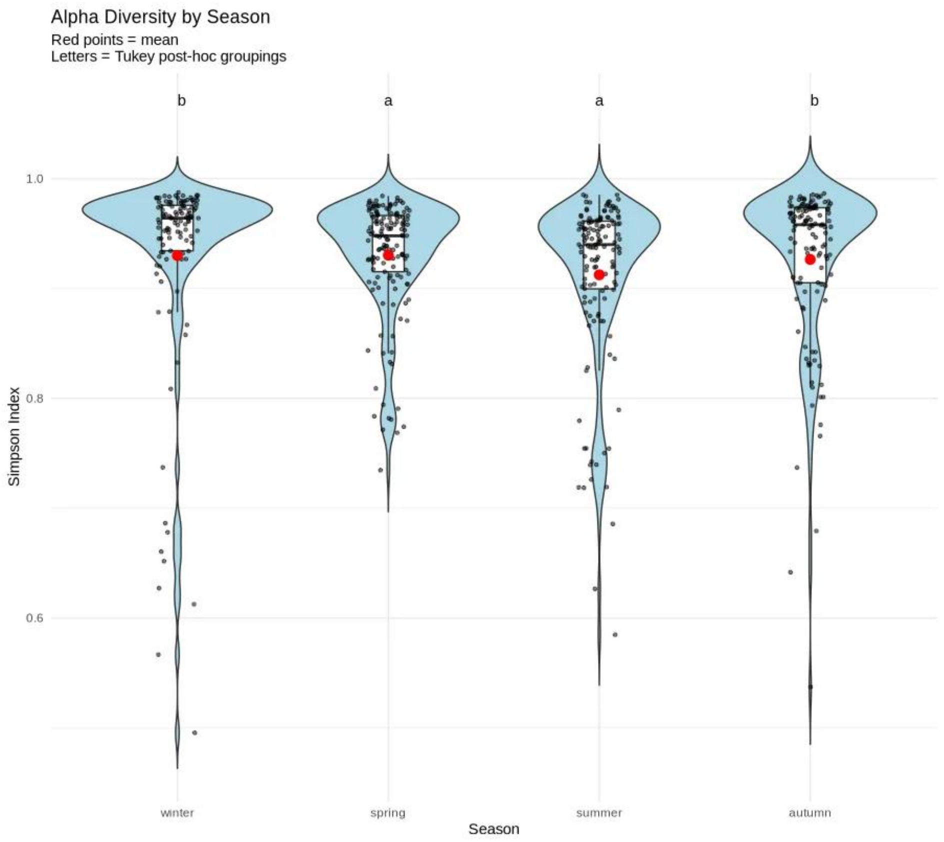
Alpha Diversity by Season
Red points = mean
Letters = Tukey post-hoc groupings
b
a
a
b
1.0
0.8
0.6
Simpson Index
winter
spring
summer
autumn
Season

**B.**

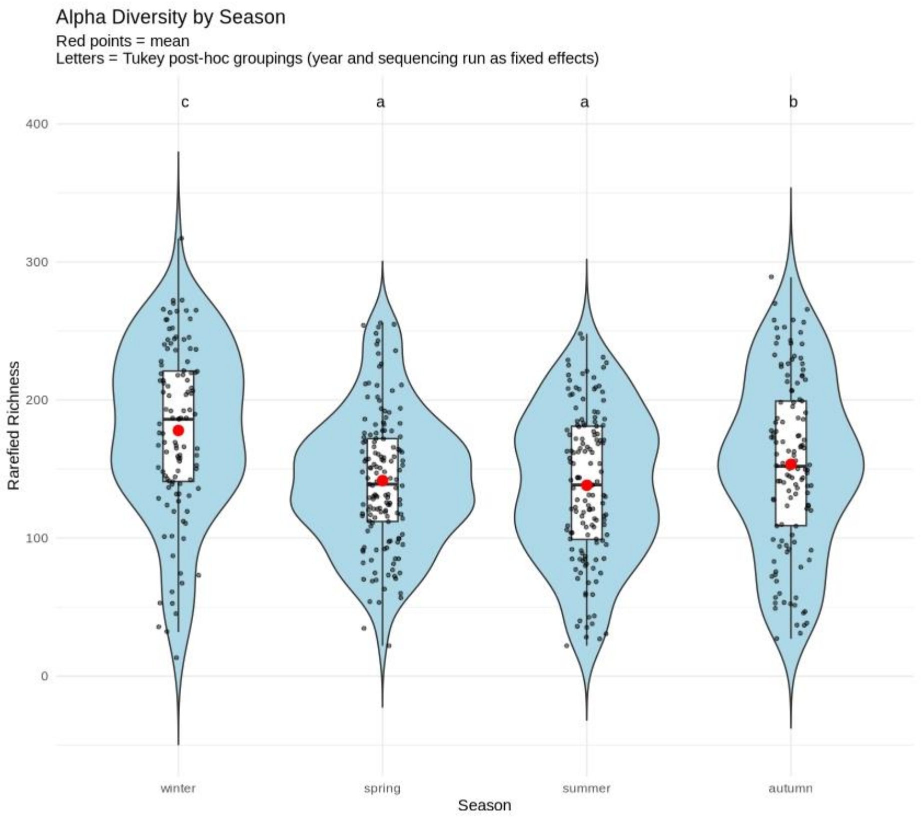


C.

**Supporting Figure 3.** Alpha diversity (A: Shannon; B: Simpson; C. Rarefied ASV Richness) by season with average means (red points); seasons with different letters were significantly different (Tukey post-hoc pairwise comparisons (Holm-adjusted) of Estimated Marginal Means: EMMs)

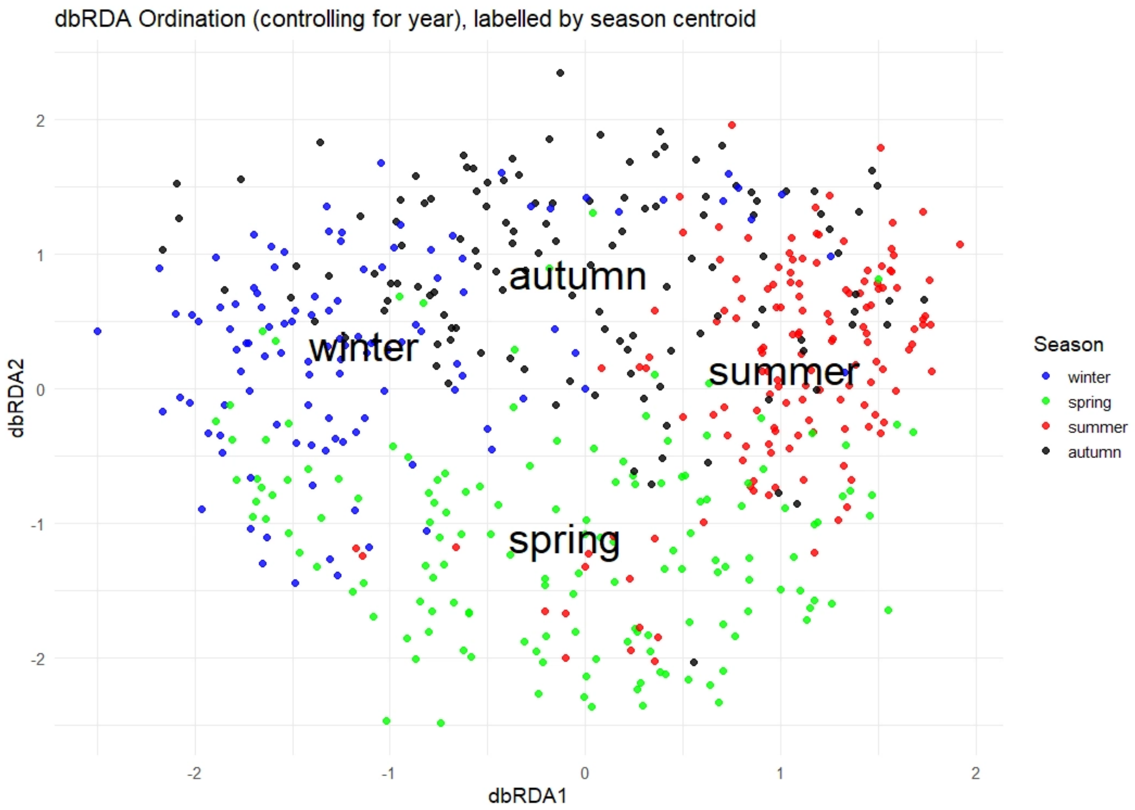


A.

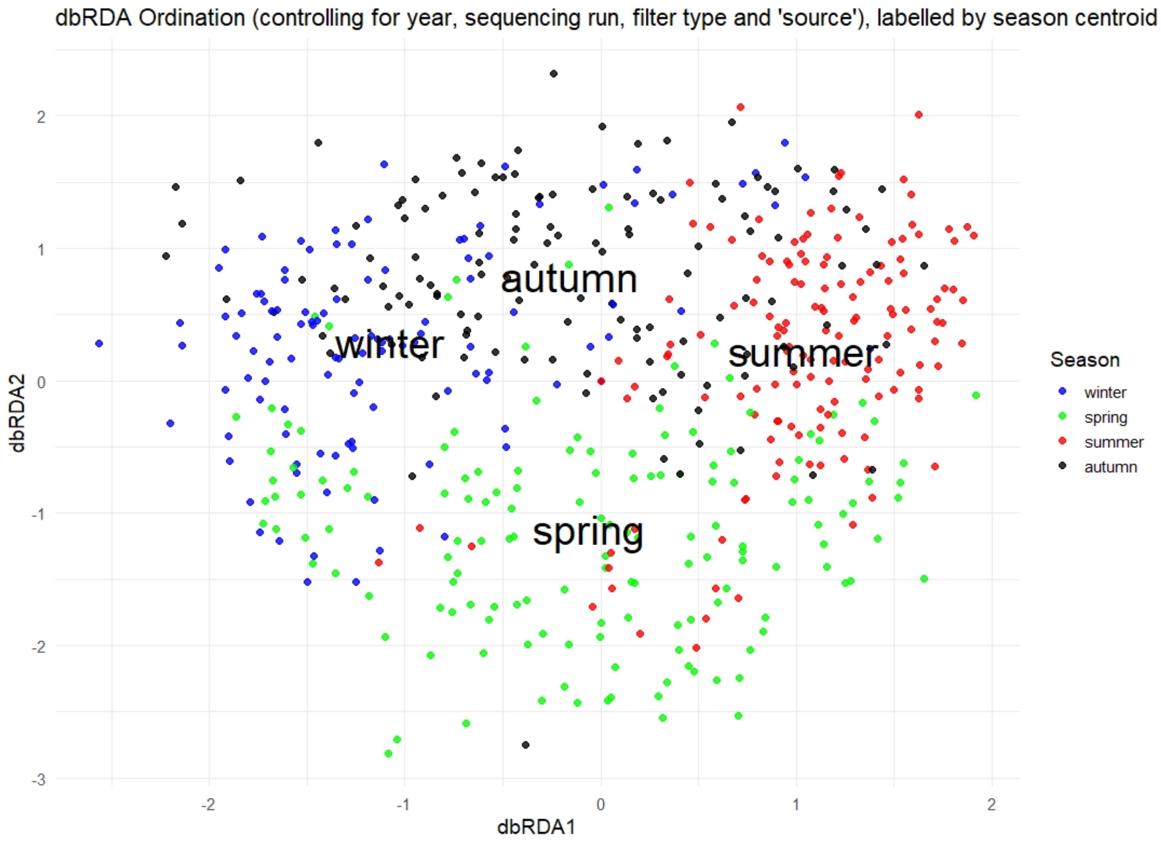


**B.**

**Supporting Figure 4.** Seasonal succession of community (ASV) composition via Distance-based Redundancy Analysis (db-RDA) models: A. Testing for the effect of season and controlling for year: B. Testing for the effect of season and controlling for year, sequencing run and inconsistency in sampling methods; points are coloured by season (Winter: blue; Spring: green; Summer: red; Autumn: black) and seasonal centroids are labelled.

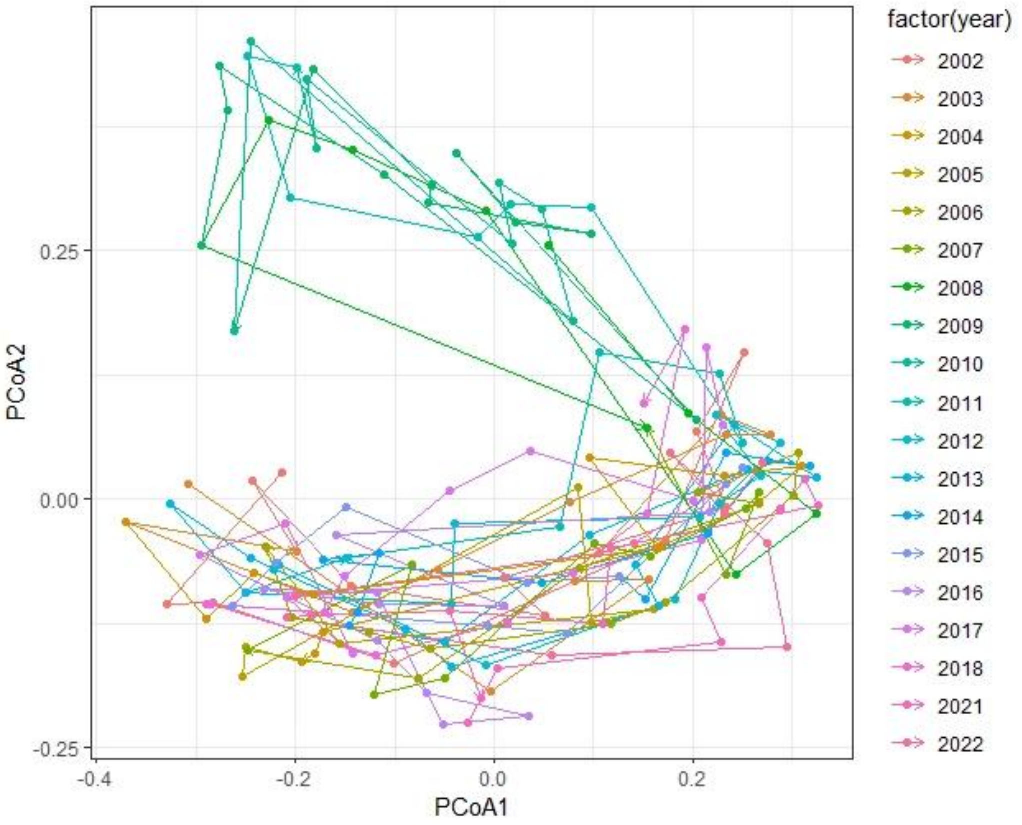


**A.**

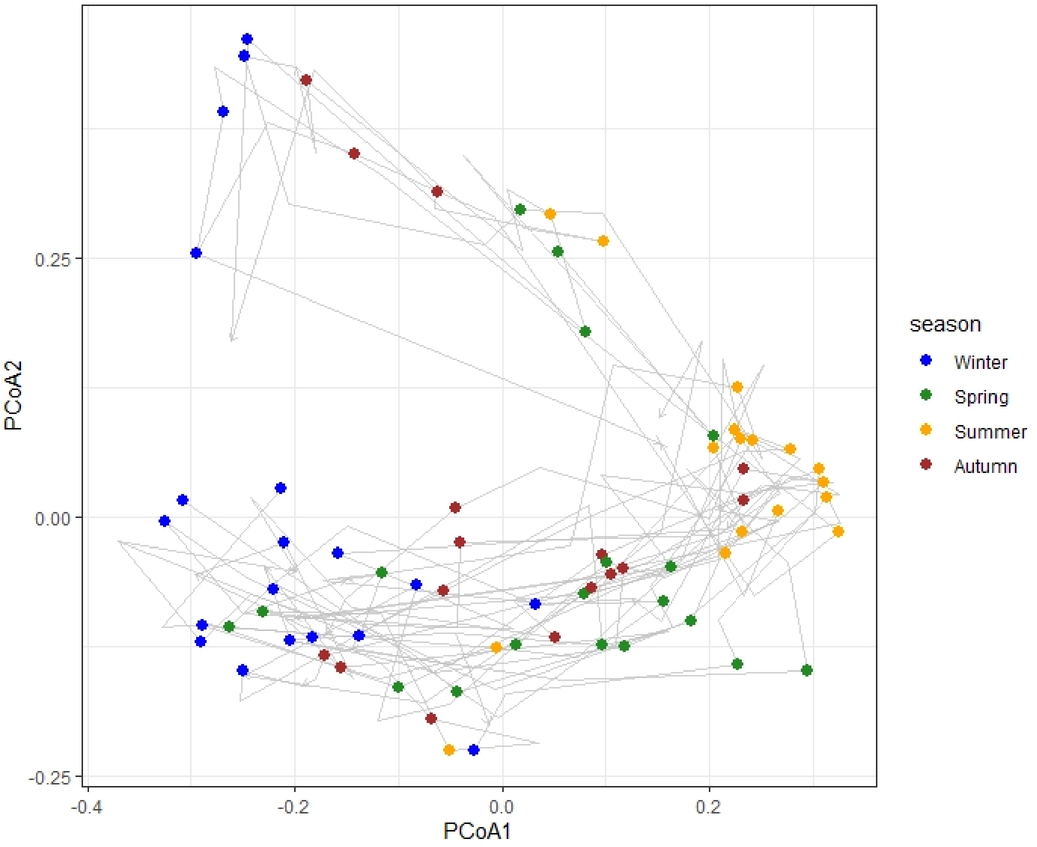


**B.**

**Supporting Figure 5.** Cyclical Ecological Trajectory Analysis (CETA) of WCO-L4 plankton community (i.e. ASVs) time series: A. Ordination of all cycles in the timeseries coloured by year; B. Ordination of cycles with points coloured by season.

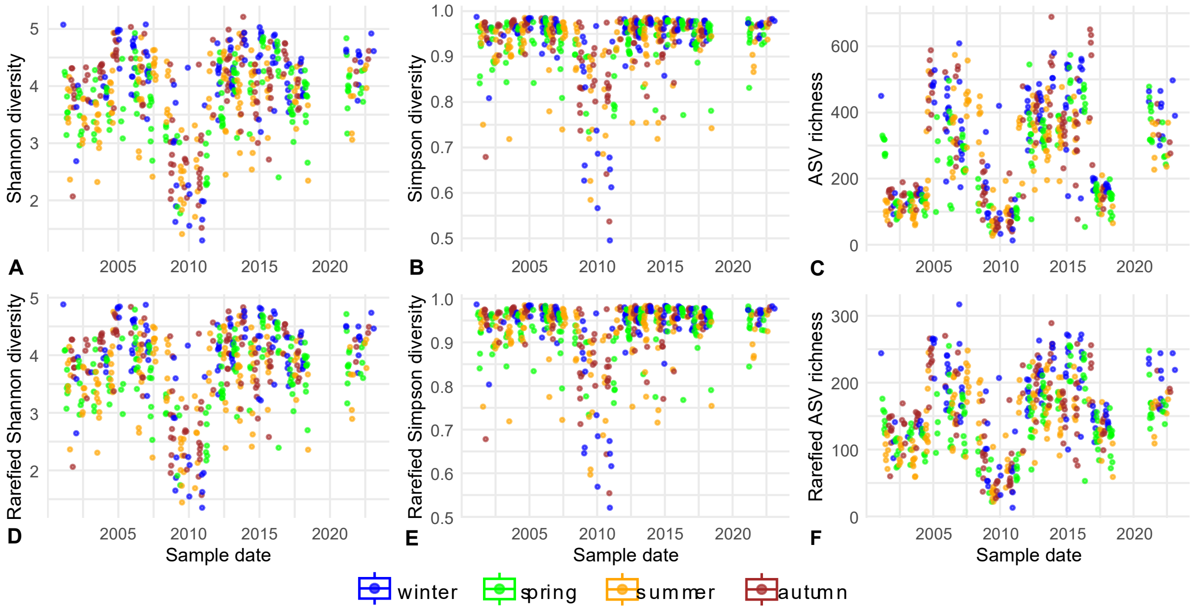


**Supporting Figure 6**. Inter-annual diversity: Per sample alpha diversity at Western Channel Observatory coastal site ‘L4’ from a long-term (years 2001-2023) eukaryote eDNA (metabarcoding: 18S-V9 rRNA) time series as calculated from the full dataset (A. Shannon; B. Simpson; C. ASV richness) and a subsampled/rarefied dataset (D. rarefied Shannon; E. rarefied Simpson; F. rarefied ASV richness); coloured by season (Winter: blue; Spring: green; Summer: yellow; Autumn: red).

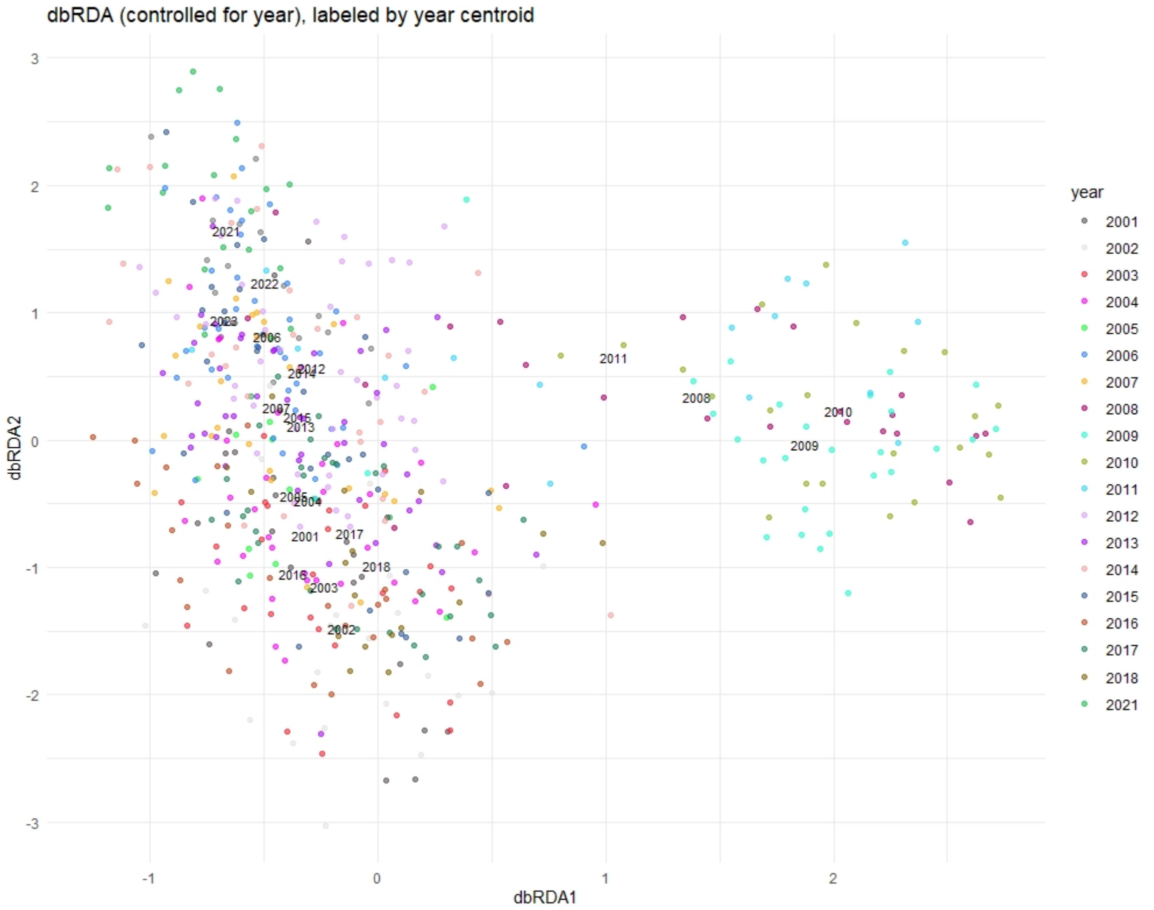

dbRDA (controlled for year), labeled by year centroid
dbRDA1
dbRDA2
year
2001
2002
2003
2004
2005
2006
2007
2008
2009
2010
2011
2012
2013
2014
2015
2016
2017
2018
2021


**A.**

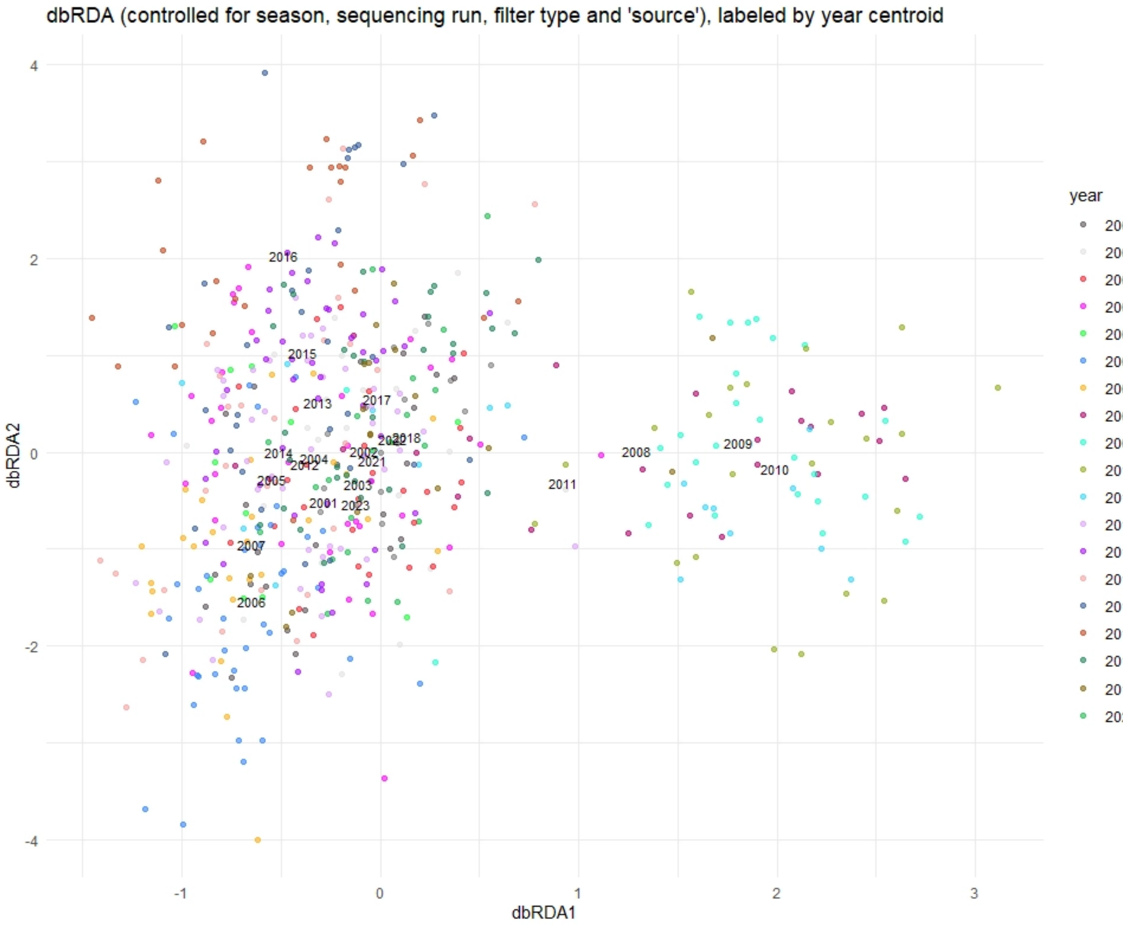


**B.**

**Supporting Figure 7.** Inter-annual eukaryote diversity at Western Channel Observatory coastal site 'L4' across years (2001-2022; eDNA metabarcoding 18S-V9 rRNA). Visualisations of community (ASV) diversity Distance-based Redundancy Analysis (db-RDA) model. A. Testing for the effect of year and controlling for season; B. Testing for the effect of year and controlling for season, sequencing run and inconsistency in sampling methods. Points are coloured by year and year centroids are labelled.

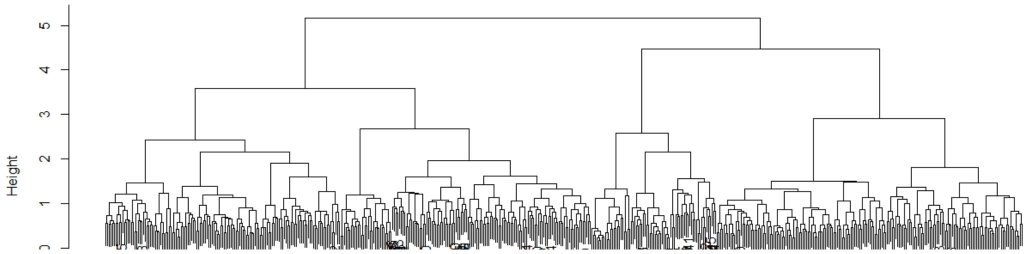


**Supporting Figure 8.** Cluster dendrogram of Hierarchical Agglomerative Clustering (HAC) of Bray-Curtis dissimilarity; Average Silhouette Width (ASW) obtained for clustering into 2-4 groups: K = 2, ASW = 0.07; K = 3, ASW = 0.07; K = 4, ASW = 0.06).

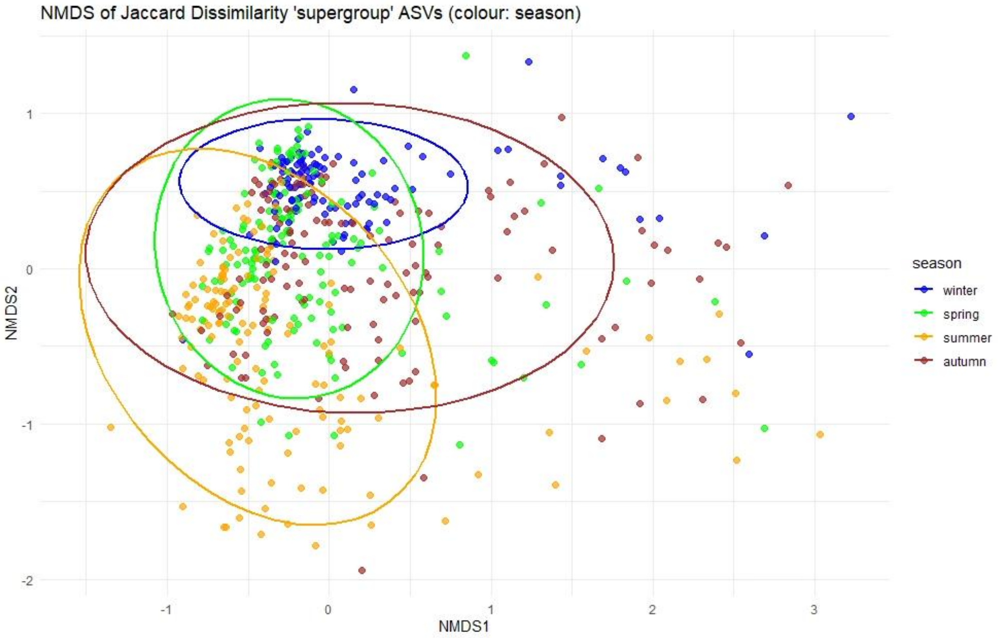


A.

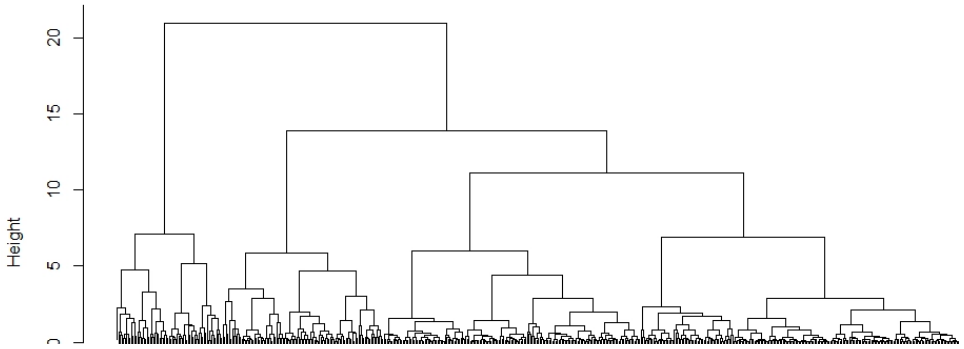


B.

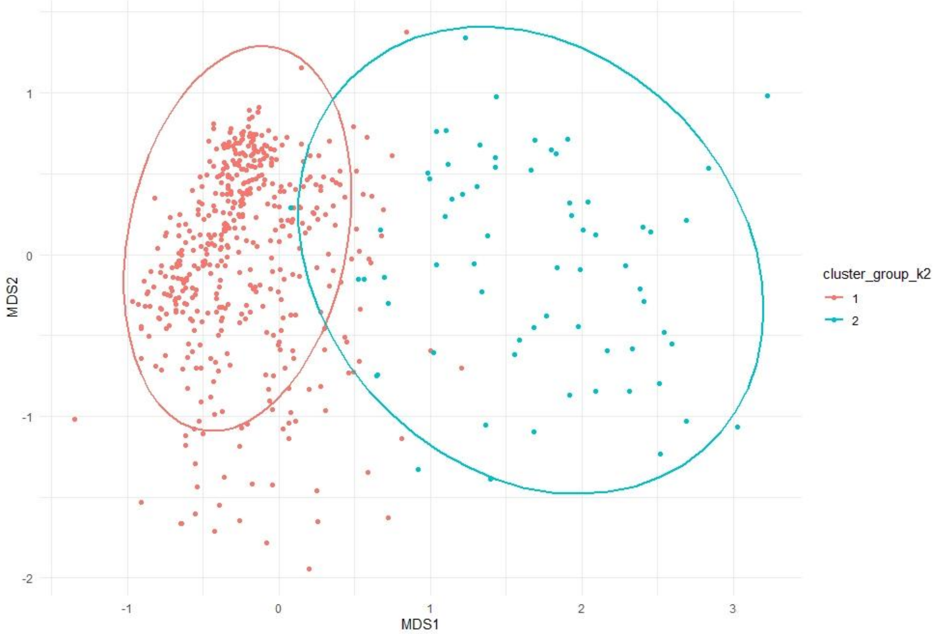


C.

**Supporting Figure 9.** Community diversity of a eukaryote metabarcoding (18S-V9 rRNA) time series from Western Channel Observatory coastal site ‘L4’ across years (2001-2023); A. NMDS ordination (K = 3) of Jaccard Dissimilarity (stress = 0.13; ADONIS test; ellipses 95% confidence interval; points coloured by season: Winter: blue; Spring: green; Summer: yellow; Autumn: red); B. Cluster dendrogram of Hierarchical Agglomerative

Clustering (HAC) of NMDS scores; C. MNDs ordination of Jaccard dissimilarity (as shown in A), with points are coloured to reflect hierarchical division into two clusters (ADONIS test; ellipses 95% confidence interval); Cluster 2 (coloured blue; right of plot) is composed entirely of samples from the years 2008-2011 (Average Silhouette Width (ASW) obtained for clustering into 2-4 groups: K = 2, ASW = 0.51; K = 3, ASW = 0.36; K = 4, ASW = 0.30).

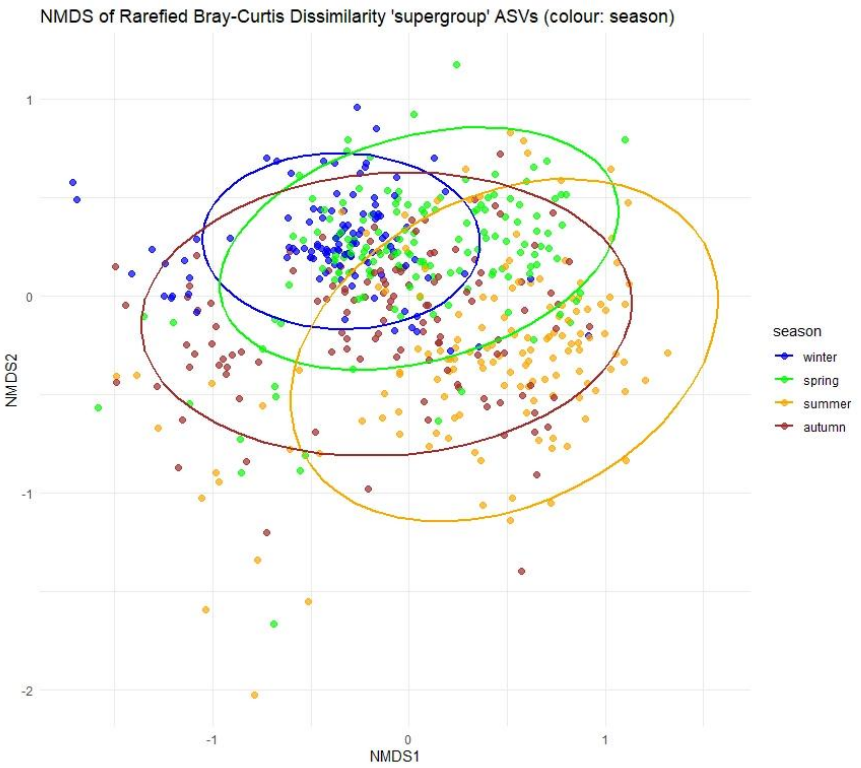


A.

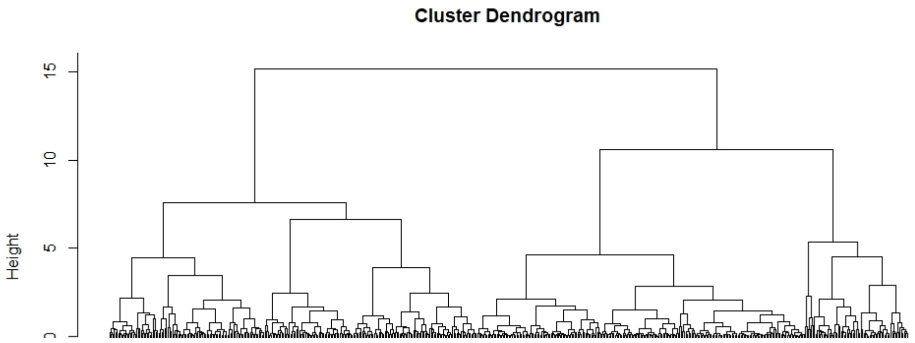


B.

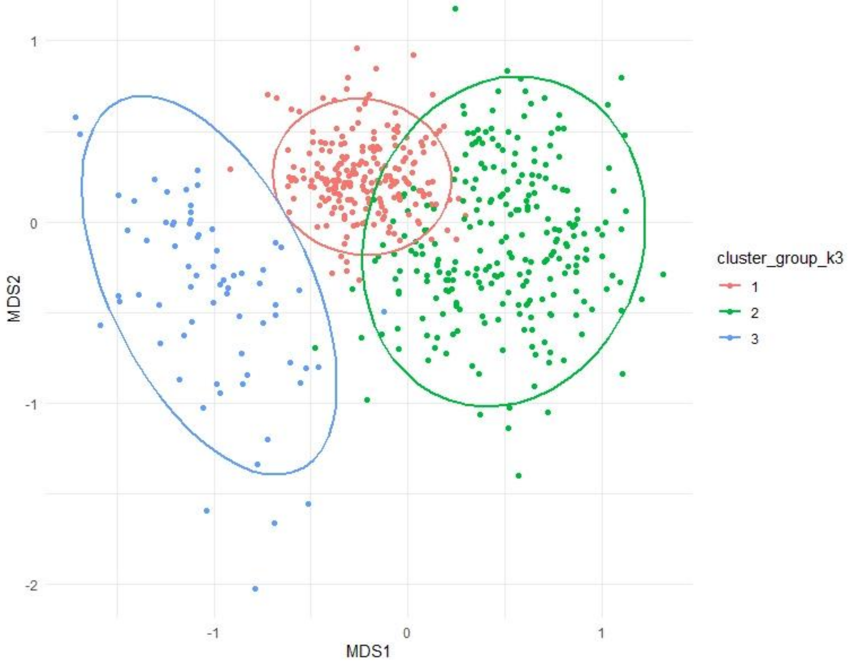


C.

**Supporting Figure 10.** Community diversity of a subsampled (i.e. rarefied to a common sequencing depth of 1145 reads) eukaryote metabarcoding (18S-V9 rRNA) time series from Western Channel Observatory coastal site ‘L4’ across years (2001-2023); A. NMDS ordination (K = 3) of Bray-Curtis Dissimilarity (stress = 0.17; ADONIS test; ellipses 95% confidence interval; points coloured by season: Winter: blue; Spring: green; Summer: yellow; Autumn: red); B. Cluster dendrogram of Hierarchical Agglomerative Clustering (HAC) of NMDS scores; C. MNDs ordination of Bray-Curtis dissimilarity (K = 3) with points are coloured to reflect hierarchical division into three clusters (ADONIS test; ellipses 95% confidence interval); Cluster 3 (coloured blue; left of plot) is composed entirely of samples from the years 2008-2011 (Average Silhouette Width (ASW) obtained for clustering into 2-4 groups: K = 2, ASW = 0.31; K = 3, ASW = 0.31; K = 4, ASW = 0.30).

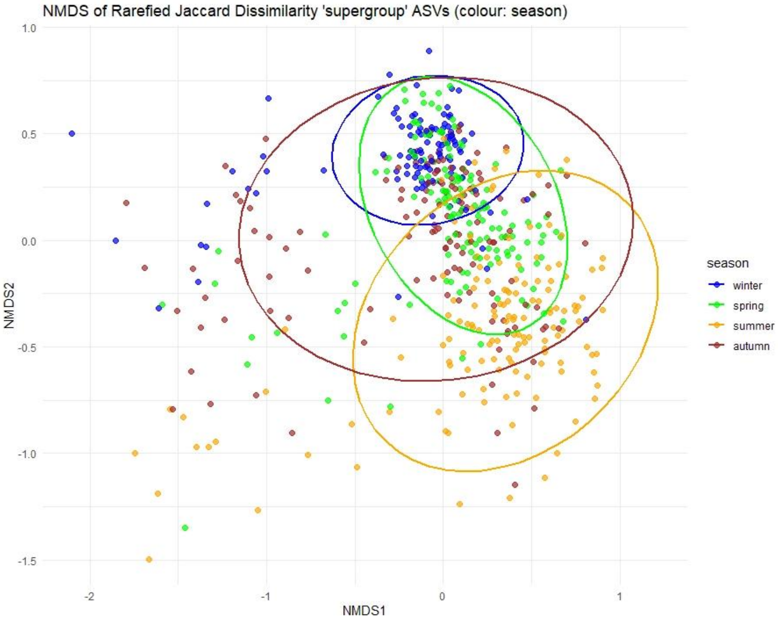


A.

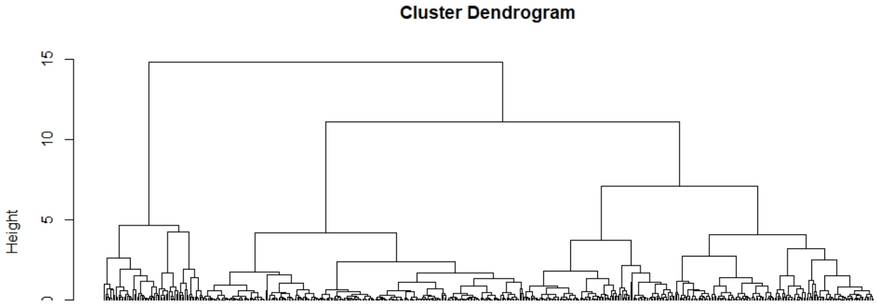


B.

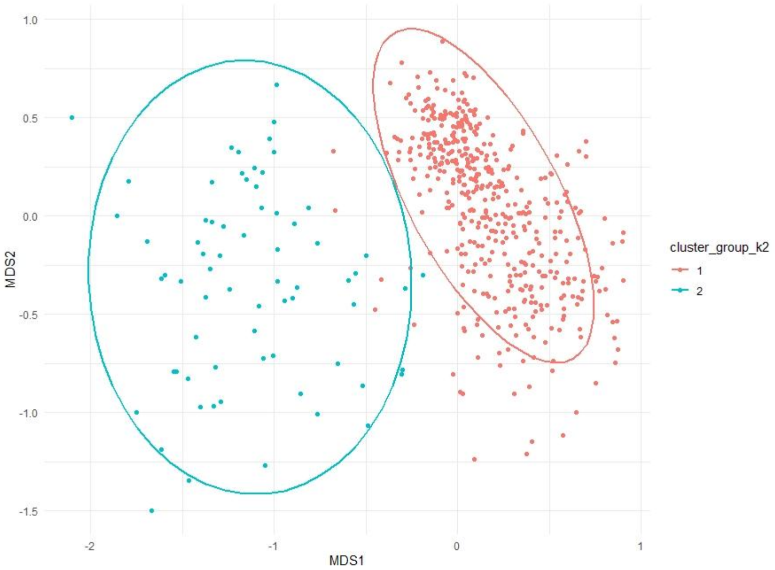


C.

**Supporting Figure 11.** Community diversity of a subsampled (i.e. rarefied to a common sequencing depth of 1145 reads) eukaryote metabarcoding (18S-V9 rRNA) time series from Western Channel Observatory coastal site ‘L4’ across years (2001-2023); A. NMDS ordination (K = 3) of Jaccard Dissimilarity (stress = 0.14; ADONIS test; ellipses 95% confidence interval; points coloured by season: Winter: blue; Spring: green; Summer: yellow; Autumn: red); B. Cluster dendrogram of Hierarchical Agglomerative Clustering (HAC) of NMDS scores; C. MNDs ordination of Jaccard dissimilarity (as shown in A), with points are coloured to reflect hierarchical division into two clusters (ADONIS test; ellipses 95% confidence interval); the majority (67/69) of samples in Cluster 2 (coloured blue; right of plot) are from the years 2008-2011 (Average Silhouette Width (ASW) obtained for clustering into 2-4 groups: K = 2, ASW = 0.50; K = 3, ASW = 0.35; K = 4, ASW = 0.34).

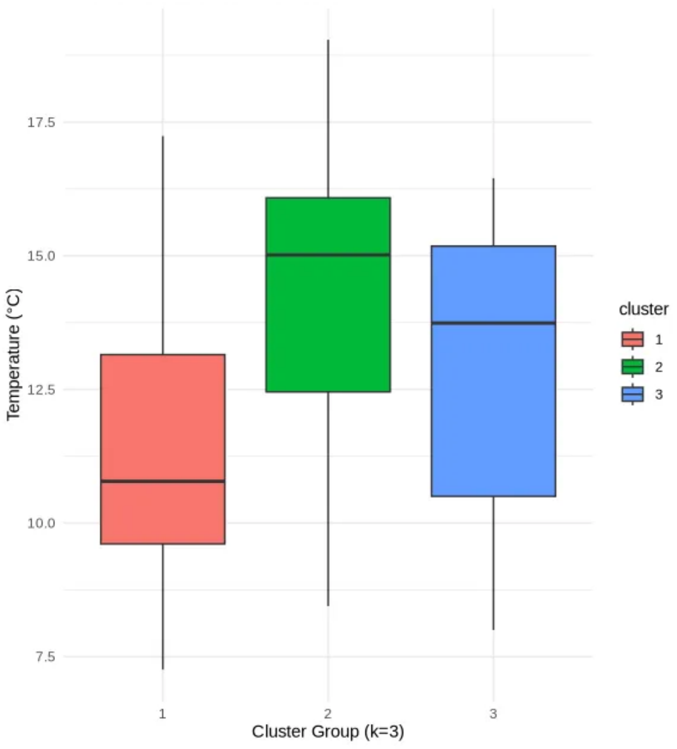
17.5
15.0
12.5
10.0
7.5
Temperature (°C)
1
2
3
Cluster Group (k=3)
cluster
1
2
3

**A.**

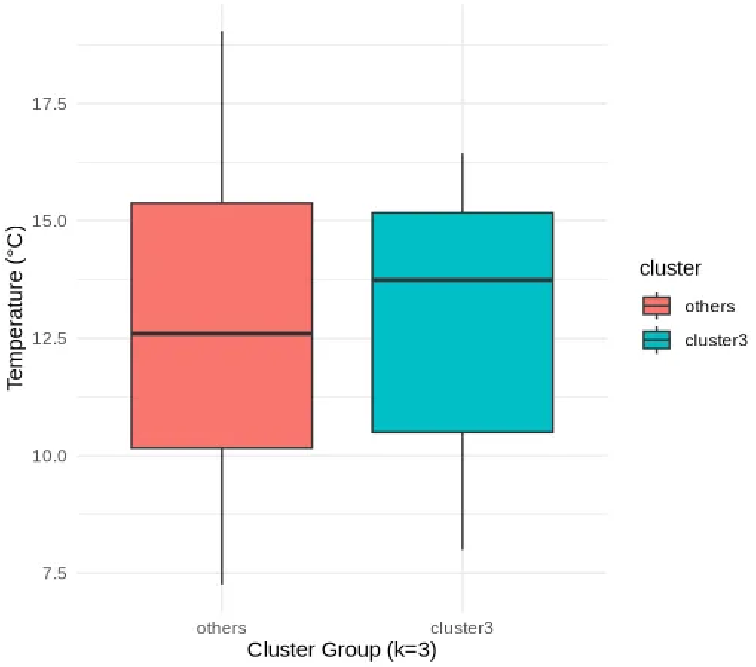
17.5
15.0
12.5
10.0
7.5
Temperature (°C)
others
cluster3
Cluster Group (k=3)
cluster
others
cluster3

**B.**

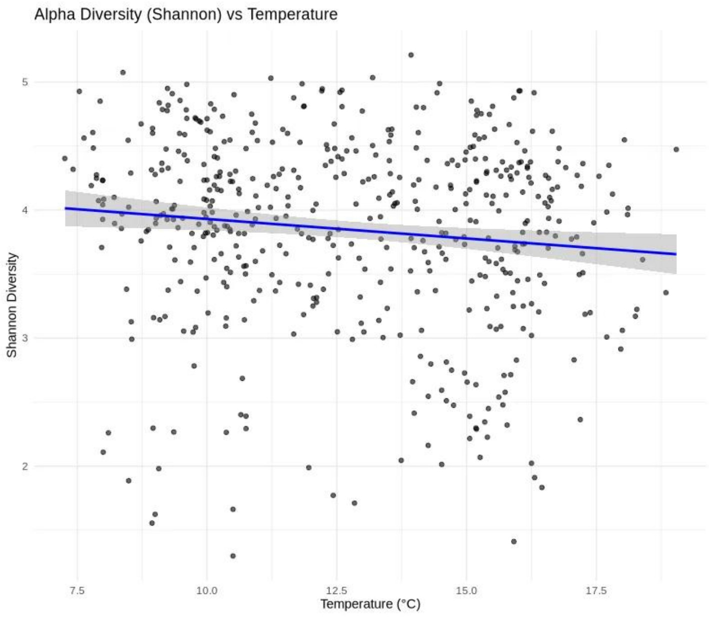


**C.**

**Supporting Figure 12**. Temperature and plankton community (ASV) diversity in the WCO-L4 plankton time series: **A.** Temperature across three groups of samples identified via hierarchical clustering; **B.** Temperature across 'Cluster 3' samples and all other samples as identified via hierarchical clustering; **C.** Correlation between temperature and Shannon Diversity (weakly negative).

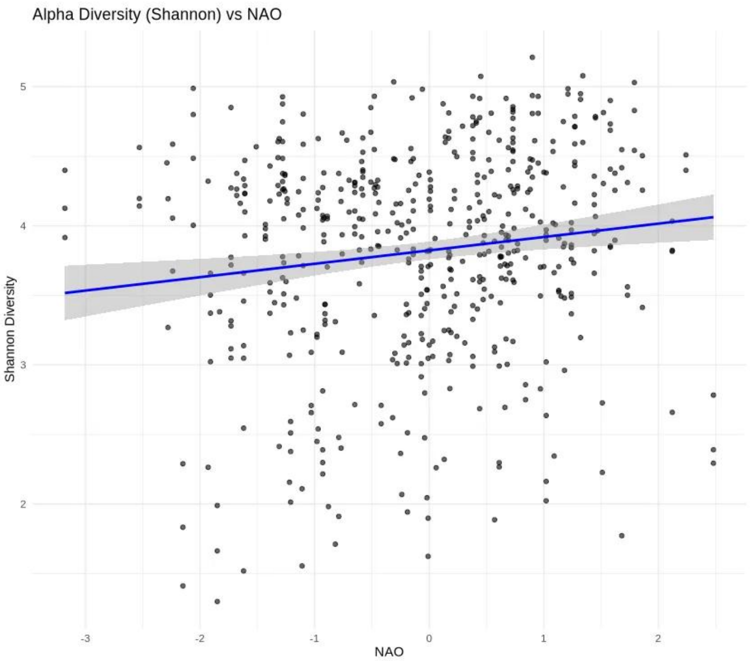


**Supporting Figure 13**. Correlation between NAO index values and Shannon Diversity (weakly positive).
